# All-optical mapping of cAMP transport reveals rules of sub-cellular localization

**DOI:** 10.1101/2023.06.27.546633

**Authors:** Katherine M. Xiang, Pojeong Park, Shon A. Koren, Rebecca Frank Hayward, Adam E. Cohen

## Abstract

Cyclic adenosine monophosphate (cAMP) is a second messenger that mediates diverse intracellular signals. Studies of cAMP transport in cells have produced wildly different results, from reports of nearly free diffusion to reports that cAMP remains localized in nanometer-scale domains. We developed an all-optical toolkit, cAMP-SITES, to locally perturb and map cAMP transport. In MDCK cells and in cultured neurons, cAMP had a diffusion coefficient of ∼130 µm^2^/s, similar to the diffusion coefficients of other small molecules in cytoplasm. In neuronal dendrites, a balance between diffusion and degradation led to cAMP domains with a length-scale of 27 ± 11 µm (mean ± s.d.). Geometrical confinement by membranes led to subcellular variations in cAMP concentration, but we found no evidence of nanoscale domains or of distinct membrane-associated and cytoplasmic pools. We discuss the scaling relations which govern diffusible signaling in tube-shaped structures.

## Introduction

The subcellular distribution of GPCR signals is thought to play a critical role in governing downstream effects (*1–3*), but the rules governing second-messenger spread are not well understood. How far and fast does the influence of local GPCR activation spread, and how is this spread affected by cell geometry and by sub-cellular localization of components of the signaling cascade? Here we develop an all-optical approach to study the sub-cellular spread of the second messenger cyclic adenosine monophosphate (cAMP), and characterize this transport in MDCK cells and neurons.

Spatial localization of cAMP signals has been proposed to play a key role in its signaling specificity (*4–8*), e.g. allowing different GPCRs to share cAMP as a second messenger, yet trigger different cellular responses (*1*, *2*, *8*). Local cAMP signaling has been demonstrated in cardiomyocytes, (*4*, *8*, *9*), in the primary cilium (*5*), and in neurons (*10*). In neurons, localized cAMP signaling may contribute to synaptic tagging (*11*), clustered plasticity (*12*), axon growth and guidance (*13*), and differential effects of neuromodulators on cell excitability and gene expression (*14*, *15*). In these examples, the cAMP confinement is essentially geometrical: impermeable membrane boundaries or long membrane-bounded tubes create localized pools which have weak diffusive coupling to other compartments.

Spatially structured enzymatic action can also shape the cAMP profile (*3*, *16–19*). Local adenylyl cyclase (AC) activity can create regional sources of cAMP, while local phosphodiesterase (PDE) activity can create regional sinks (*20*). Distinct distributions of sources and sinks can create nonequilibrium steady-state concentration profiles (*21*). These gradients have been observed in neurons (*22*, *23*), migrating cells (*24*), and cardiomyocytes (*25*). Experiments comparing membrane-targeted vs. cytoplasmic cAMP reporters in HEK cells found cAMP gradients between membrane and cytosol upon stimulation of plasma membrane GPCRs, (*26*, *27*) attributed to differential action of membrane-bound vs. cytosolic PDEs (*28*). However, the spatial profile of the signaling was not resolved in these experiments.

Recent work claimed that nanometer-sized cAMP-depleted domains exist around individual PDE molecules (*29–31*), and cAMP-enriched domains around individual GPCR molecules (*32*) (for reviews, see (*33*, *34*)). These works suggested that locally scaffolded “signalosomes” could link sensor, transducer, and effector proteins in a macromolecular complex, governed by a private pool of cAMP. However, based on the data available, it is not clear to what extent the cAMP is localized around individual molecular sources, or if the clustering exists within higher-order structures which share cAMP among ensembles of sources and receptors. Other recent work suggested that phase-separated liquid droplets could also act as hubs for private cAMP signaling (*35*).

A challenge for models of nano-localized cAMP signaling is that these ideas are inconsistent with simple models based on long-established understanding of small-molecule transport and enzyme kinetics (*21*, *36–40*) (reviewed in (*41*)). A scaling argument illustrates the problem:

Coupled diffusion and homogeneous degradation (e.g. by PDEs) lead to a length-scale (the “Thiele length”), corresponding to the mean distance a cAMP molecule diffuses before it is degraded (*42*, *43*). This sets the size of the signaling domain. The Thiele length in homogeneous solution is 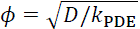, where *D* (μm^2^/s) is the cAMP diffusion coefficient and *k*_PDE_ (s^-1^) is the first-order cAMP degradation rate (equal to *k*_cat_[PDE]/*K*_M_, provided that [cAMP] « *K*_M_) ^1^. Applying reasonable values for diffusion of small molecules in cytoplasm (*D* = 50 – 500 μm^2^/s) (*44*, *45*) and the kinetics of PDE activity in neurons (*k*_PDE_ = 0.12 – 6.15 s^-1^ for [PDE] = 0.4 µM) (*46*, *38*, *41*) gives ф = 3 — 65 μm, comparable to the size of most cells. For the Thiele length to decrease to single-nanometer dimensions (a factor of ∼10^4^) would require an increase in *k*_PDE_ by 10^4^ *and* a decrease in *D* by 10^4^, neither of which is plausible.

A similar argument based on spherical diffusion yields the deviation in concentration near a single-molecule sink or source (see **Supplemental Materials Section 1** for details). At a distance of 1 nm from an enzyme, the concentration change (in nM) is Δ*c*(1 nm) ≈ ±132 *k*_cat_/*D*, where *k*_cat_ (s^-1^) is the enzyme turnover rate, *D* (μm^2^/s) is the cAMP diffusion coefficient and the sign depends on whether the enzyme is a source (+) or sink (-). For a saturating turnover rate of *k*_cat_ = 5 s^-1^ (corresponding to PDE4 (*41*)) and a diffusion coefficient *D* = 100 μm^2^/s, Δ*c*(1 nm) = —6.6 nM. On a baseline cAMP concentration of ∼1 μM (*41*, *47*), this deviation is negligible. For the local deviation to approach the baseline level of cAMP would require an increase in the ratio *k*_cat_/*D* by a factor of ∼100.

Cyclic AMP binds to intracellular buffers, such as protein kinase A (PKA), which can slow its diffusion. The influence of “buffered diffusion” has been studied in detail for Ca^2+^ (*42*, *48–50*), and similar principles apply to cAMP (*30*). Reversible binding to an immobile buffer decreases *D*; but also protects cAMP from enzymatic degradation and hence decreases *k*_PDE_ proportionally. These effects cancel in the expression for 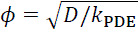. Hence, the presence of the buffer does not affect the Thiele length (*42*, *43*, *51*). An immobile buffer also does not affect the concentration profile of free cAMP around a single-molecule source or sink.

While the estimates above present a challenge to claims of nanoscale localization, the underlying assumptions may not apply in cells. Nonlinear interactions, such as cAMP-dependent PDE activation via phosphorylation (*52*, *53*), bidirectional interactions between cAMP and Ca^2+^ signaling pathways (*54–56*), or multimeric enzyme assemblies (*21*) might affect cAMP profiles in unexpected ways.

Motivated by the dramatic discrepancy between recent reports of nanoscale localization and our simple biophysical estimates, we sought to characterize cAMP transport in cells experimentally. We developed bicistronic constructs to co-express a blue light-activated adenylyl cyclase, bPAC (*57*), with a yellow-excited fluorescent cAMP reporter, Pink Flamindo (PF) (*58*), with distinct sub-cellular localization tags on the two proteins. We call these tools “cAMP Sub-cellular Indicators of Transport”, or cAMP-SITES.

By combining sub-cellular molecular and optical targeting, we probed cAMP transport over length-scales from nanometers to ∼100 µm. We found that cAMP diffused relatively freely in MDCK cells and in neurons, with a diffusion coefficient of 130 µm^2^/s. Neuronal dendrites supported cAMP “domains” of ∼25 µm, set by the balance of diffusion and degradation. We found no evidence of nanoscale domains. Cyclic AMP concentrations were strongly affected by local surface-to-volume ratio, and by whether production and degradation each occurred in solution or on the membrane. We introduce scaling rules which relate cell geometry to the localization and spread of soluble signals, and compare to the spread of electrical signals.

## Results

### An all-optical toolkit for studying cyclic AMP transport

To enable independent optical perturbation and measurement of cAMP in cells, we first created the cAMP-SITE construct CMV::PinkFlamindo-p2a-bPAC (hereafter, PF+bPAC), where p2a is a self-cleaving peptide linker (Fig. 1A). Blue (488 nm) light activated the bPAC via an endogenous flavin chromophore to drive cAMP production. Yellow (561 nm) light excited Pink Flamindo (PF) fluorescence to probe the cAMP dynamics. To achieve sub-cellular control of bPAC activation, we used a digital micromirror device (DMD) to pattern the blue light with micron-scale precision (Fig. 1B, Methods). After a brief (0.5 s) pulse of patterned blue light delivered to a field of HEK293T cells, PF fluorescence increased in the illuminated regions over ∼40 s and then returned to baseline over several minutes (Fig. 1C). Similar dynamics were observed when bPAC was activated in subcellular domains in MDCK cells (Fig. 1C). In this case the cAMP signal gradually diffused out of the directly stimulated regions but remained within the contours of the stimulated cells.

**Figure 1.**
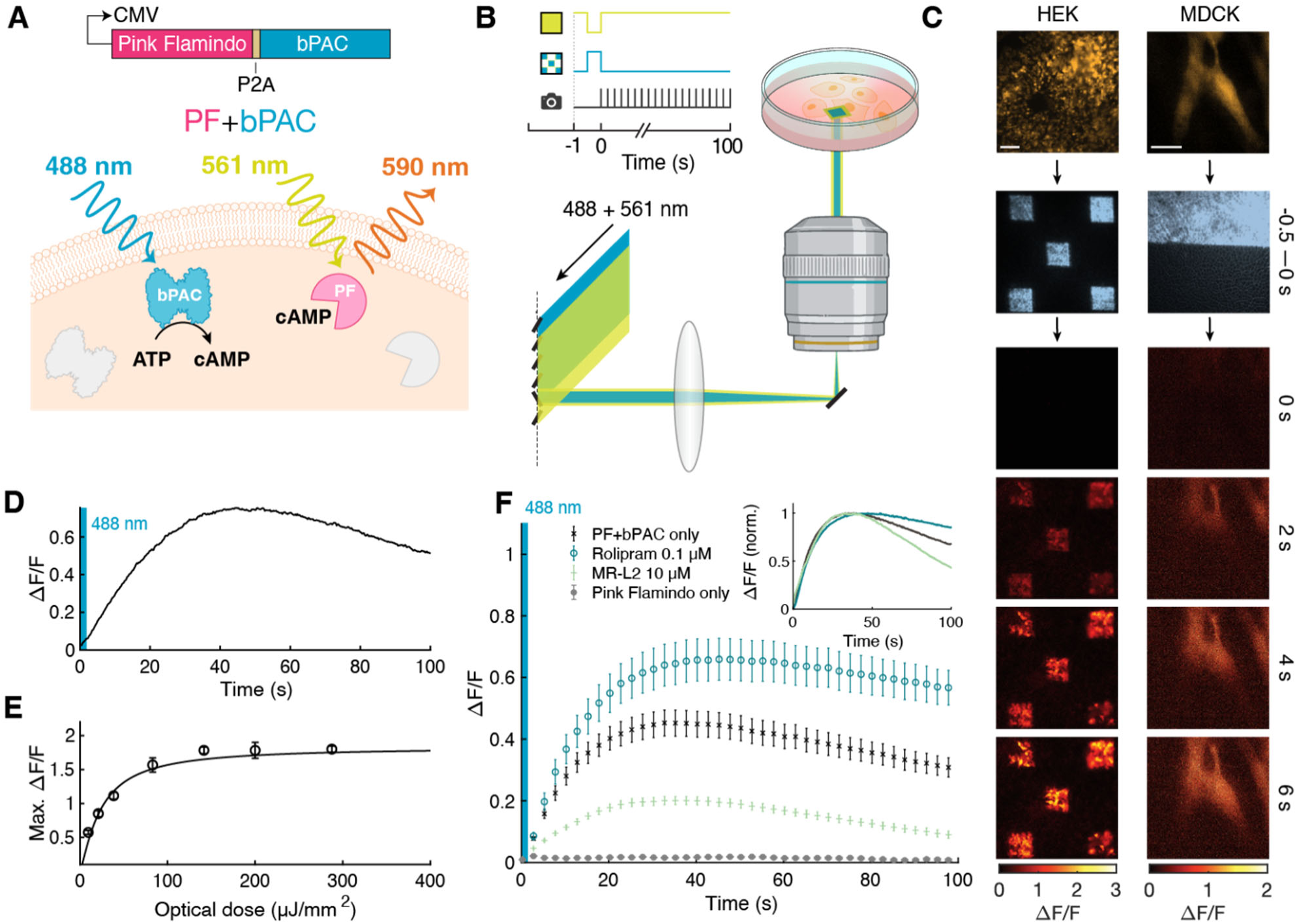
Targeted optogenetic production and measurement of cAMP. **A)** Genetic construct for optogenetic production and imaging of cAMP. **B)** Illumination protocol for patterned optogenetic activation of bPAC (blue, 488 nm, 0.5 s), followed by wide-field imaging of PF (yellow, 561 nm). **C)** Top: basal PF fluorescence in a dish of HEK cells (left, scale bar: 100 µm) and MDCK cells (right, scale bar: 20 µm) expressing PF+bPAC. Middle: Patterns of transient (0.5 s) blue illumination for bPAC activation. Bottom: images of PF ΔF/F at 2-s intervals show subsequent cAMP dynamics. Note intracellular spread of cAMP away from the blue-illuminated region in the MDCK cells. **D)** Example single-cell ΔF/F trace in a HEK cell after 0.5 s wide-field blue illumination. **E)** Blue light dose-response curve of PF+bPAC, measured by peak ΔF/F. Fit to Hill equation gives saturating ΔF/F = 1.8 ± 0.2 (mean ± s.d.), half-saturating blue light intensity 24 ± 5 µJ/mm^2^, and Hill coefficient 1.2 ± 0.3 (95% CI from the computed Jacobian, *n* = 11 – 60 cells per data-point. Error bars represent SEM.) **F)** Response of HEK cells expressing PF+bPAC subject to the protocol in (D) with 21 µJ/mm^2^ blue light (black, *n* = 40 cells from 3 dishes). Cells treated with the PDE inhibitor rolipram (blue, *n* = 19 cells from 2 dishes) had slower recovery and larger peak ΔF/F. Cells treated with the PDE activator MR-L2 (green, *n* = 39 cells from 2 dishes) had faster recovery and smaller peak ΔF/F. Cells expressing PF without bPAC (gray, *n* = 42 cells from 2 dishes) did not respond to blue light. Inset: peak-normalized traces. Error bars represent SEM.

As blue light dose increased, the peak PF fluorescence response increased linearly and then saturated at ΔF/F = 1.8 ± 0.2 (*n* = 38 HEK cells, mean ± s.d., Fig. 1D–E), with a half-maximal response at a blue illumination dose of *I*_1/2_ = 24 ± 5 µJ/mm^2^. The relation between optical dose and cAMP production depends on the expression level of bPAC and on phosphodiesterase (PDE) activity, so the value of *I*_1/2_ could vary between cell types and experimental conditions. Treatment with 100 µM forskolin, a broad-spectrum AC activator, and 200 µM IBMX (3-isobutyl-1-methylxanthine, a broad-spectrum PDE inhibitor) did not increase the PF ΔF/F beyond the level induced by saturating blue light (Fig. S1A), implying that the saturation behavior observed in Fig. 1E was likely PF saturation. To avoid saturating the PF reporter, we adjusted blue light doses to keep ΔF/F < 1 for subsequent experiments.

We performed pharmacological tests to confirm that PF reported cAMP (Table 1). In HEK cells expressing PF+bPAC, rolipram, a PDE4 inhibitor, increased the magnitude and duration of the PF response to a blue light pulse (Fig. 1F). In separate experiments, MR-L2, a PDE4 activator, decreased the magnitude and duration of the PF response. Control experiments in HEK cells expressing just PF showed no response to blue excitation (Fig. 1F), confirming that the blue light did not disturb the PF mApple chromophore or perturb cAMP through pathways independent of bPAC. HEK cells expressing PF+bPAC but exposed only to steps of yellow light did not show cAMP transients, confirming that the yellow imaging light did not drive spurious bPAC activation (Fig. S1B). Together, these experiments established that the PF+bPAC pair enabled crosstalk-free all-optical perturbation and measurement of cAMP dynamics in cells.

**Table 1.** Experimental details for HEK293T cell pharmacology.

| Condition | Number of cells | Number of exp. replicates |
| --- | --- | --- |
| PF+bPAC | 40 | 3 |
| + 0.1 $\mu\text{M}$ Rolipram | 19 | 2 |
| + 10 $\mu\text{M}$ MR-L2 | 39 | 2 |
| PF only | 42 | 2 |

We then produced versions of the cAMP-SITES where one or both of the bPAC and the PF were targeted to the membrane. The goals were (a) to minimize diffusion of the source (bPAC) and reporter (PF) as confounding variables in measurements of cAMP transport, (b) to investigate the possibility of distinct cAMP pools at the membrane vs. in the cytoplasm, and (c) to develop a toolkit to explore the interaction of source localization and cell geometry on cAMP transport and dynamics. We generated four cAMP-SITES to co-express all combinations of soluble and membrane bound PF and bPAC (Fig. 2A–D, Fig. S2) and designated these: PF+bPAC, PF+bPAC^m^, PF^m^+bPAC, and PF^m^+bPAC^m^ (Table 2). The p2a linker left 21 amino acids on the C-terminus of the first protein in each sequence (*59*) and was incompatible with the also C-terminal CAAX membrane-targeting motif. Hence, we used the CAAX motif to target bPAC to the membrane and an N-terminal myristoylation motif (*60*) to target PF to the membrane (Methods).

**Figure 2.**
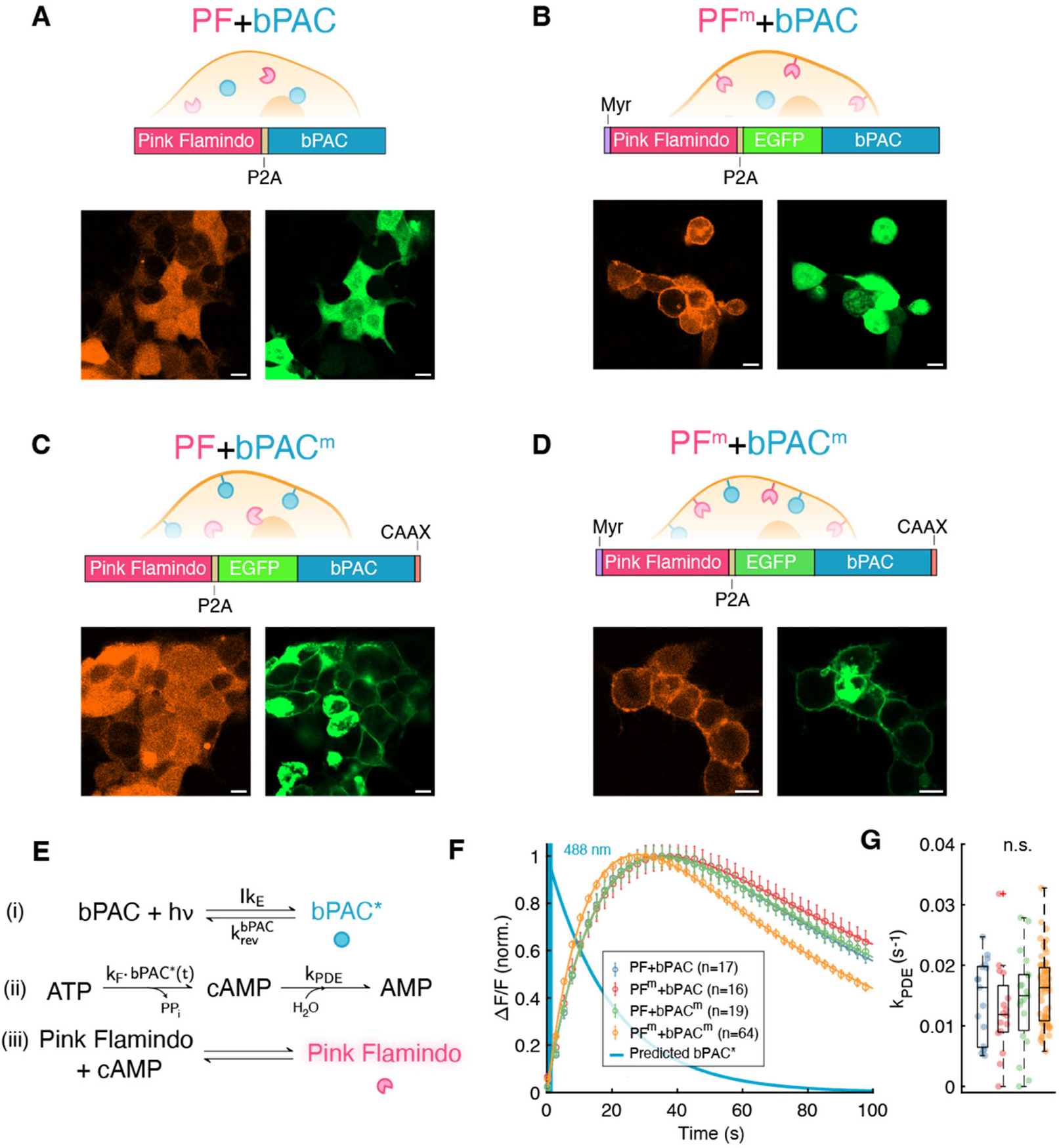
cAMP dynamics with soluble and membrane bound actuators and reporters. **A-D)** Genetic constructs for spatial control of actuator and reporter. **A)** Soluble PF and soluble bPAC. Below: single-plane confocal images of HEK cells with PF (left) and GFP-tagged bPAC (right). Scale bar: 10 µm. **B–D)** Same as **A**, but for membrane PF (PF^m^) + bPAC, PF + membrane bPAC (bPAC^m^), and for PF^m^ + bPAC^m^. **E)** Kinetic model for (i) photoactivation of bPAC, (ii) cAMP production and degradation, and (iii) cAMP sensing. Here *k*_PDE_ = *k*_cat_[PDE]/*k*_M_. **F)** Mean fluorescence responses of HEK cells expressing each combination of soluble or membrane bound bPAC and PF, with best-fit model traces overlaid (Equation 5 in **Supplemental Materials**, *R^2^*= 0.9973, 0.9988, 0.9981, 0.9982, from legend top to bottom). All traces normalized to peak. Single-cell traces were fit with *k*^bPAC^fixed within each construct and *k*_PDE_allowed to vary between cells. Error bars represent SEM. Predicted blue light impulse response (**Supplemental Materials Section 3.1**) for activated bPAC (bPAC*, blue), with *k*^bPAC^ = 0.05 s^-1^, the best-fit rate for PF+bPAC. **G)** Extracted PDE rate constants for (left to right): PF+bPAC, PF^m^+bPAC, PF+bPAC^m^, and PF^m^+bPAC^m^ (see Table 3). Boxes represent the interquartile range with the median denoted by the central line. All four conditions are not significantly different according to the Wilcoxon rank-sum test.

**Table 2.** Genetic constructs in this study.

| Construct name | Full construct | Reference |
| --- | --- | --- |
| PF+bPAC | pLenti-CMV-Pink Flamindo-p2a-bPAC | Addgene #202716 |
| PF<sup>m</sup>+bPAC | pLenti-CMV-Myr-Pink Flamindo-p2a-GFP-bPAC | Addgene #202717 |
| PF+bPAC<sup>m</sup> | pLenti-CMV-Pink Flamindo-p2a-paGFP-bPAC-CAAX | Addgene #202718 |
| PF<sup>m</sup>+bPAC<sup>m</sup> | pLenti-CMV-Myr-Pink Flamindo-p2a-GFP-bPAC-CAAX | Addgene #202719 |
| PF | Pink Flamindo | Addgene #102356 |
| bPAC | bPAC | Addgene #85468 |
| PF<sup>m</sup> | pLenti-CMV-Myr-Pink Flamindo | Addgene #204669 |
| paGFP-bPAC | pLenti-CMV-paGFP-bPAC | Addgene #204670 |
| paGFP-bPAC<sup>m</sup> | pLenti-CMV-paGFP-bPAC-CAAX | Addgene #204671 |

**Table 3.** Kinetic parameters of cAMP-SITE constructs in HEK293T cells.

| Construct | $k_{PDE}$ (s<sup>-1</sup>) | $k_{rev}^{bPAC}$ (s<sup>-1</sup>) | Number of cells | Number of exp. replicates |
| --- | --- | --- | --- | --- |
| PF+bPAC | 0.014 ± 0.007 | 0.050 ± 0.017 | 17 | 3 |
| PF<sup>m</sup>+bPAC | 0.013 ± 0.008 | 0.051 ± 0.023 | 16 | 2 |
| PF+bPAC<sup>m</sup> | 0.014 ± 0.008 | 0.050 ± 0.023 | 19 | 2 |
| PF<sup>m</sup>+bPAC<sup>m</sup> | 0.016 ± 0.006 | 0.069 ± 0.018 | 64 | 2 |

We confirmed that PF^m^ was membrane bound by directly imaging its fluorescence, and we confirmed that GFP-bPAC^m^ was membrane bound via images of GFP fluorescence (Fig. 2A–D and S2 for expression in HEK cells, Fig. S3 for neurons). While some cells expressing PF^m^ or bPAC^m^ showed intracellular aggregates, for measures of cAMP transport we selected cells with clean membrane trafficking. HEK cells expressing PF^m^ or PF showed similar forskolin dose-dependent increases in fluorescence, confirming that the membrane-targeted reporter remained functional (Fig. S4A). HEK cells co-expressing PF and either bPAC or bPAC^m^ showed similar blue light dose-dependent elevations in PF fluorescence, confirming that bPAC^m^ was also functional (Fig. S4B).

To probe the lateral diffusion of bPAC^m^, we expressed fusions of photoactivatable GFP (paGFP) to either bPAC or bPAC^m^ and used patterned illumination at 405 nm to activate the paGFP in sub-cellular regions. The soluble paGFP-bPAC spread freely while the paGFP-bPAC^m^ showed little diffusion over 100 s, confirming that the membrane-bound bPAC^m^ construct was effectively immobile on the timescale of our experiments (Fig. S5, Methods).

Prior studies had reported that bPAC has dark-state adenylyl cyclase activity, which might elevate basal cAMP (*61*, *62*). We reasoned that elevated basal cAMP would decrease the PF response to either strong bPAC activation or saturating forskolin. In cells co-transfected with PF and GFP-bPAC^m^, the bPAC^m^ expression (as quantified by GFP fluorescence) varied over ∼3 orders of magnitude. The PF response to a strong pulse of blue light (234 μJ/mm^2^) showed a bell-shaped dependence on bPAC^m^ expression (Fig. S6). On the low bPAC^m^ side, cAMP production was an increasing function of bPAC^m^ expression, as expected for the blue light-activated AC activity. On the high bPAC^m^ side, the PF response was a decreasing function of bPAC^m^ expression, consistent with dark-state bPAC^m^ activity. A simple model comprising a saturable PF response and bPAC^m^ dark-state activity (**Supplemental Materials Section 2**) reproduced this bell-shaped curve (Fig. S6B).

We then tested the response of PF^m^ to saturating (100 μM) forskolin, in the absence of blue light. At low bPAC^m^ expression, the PF^m^ response was independent of bPAC^m^; at high bPAC^m^, the response was a decreasing function of bPAC^m^ (Fig. S6C), consistent with dark-state bPAC^m^ activity. From these data, we established an empirical expression level threshold (50^th^ percentile) below which the influence of bPAC^m^ dark-state activity was minimal (Fig. S7). We also tested biPAC^m^, a photoactivated adenlylyl cyclase reported to have less dark activity (*63*), and found that its dark activity was similar to that of bPAC under our conditions (Fig. S7C). Thus, we did not pursue biPAC further.

To test the effect of dark-state activity of bPAC^m^ in neurons, we measured the ΔF/F_max_ when treated with 100 µM forskolin for neurons expressing only PF^m^ vs. neurons co-transfected with PF^m^ and bPAC^m^ (Fig. S8A). The two groups responded similarly, with ΔF/F ≈ 1.4, confirming that at the expression levels of our experiments, dark-state activity of bPAC^m^ in neurons was negligible.

Finally, we tested whether PF and PF^m^ were sensitive within the physiological range of cAMP concentrations in neurons. Both reporters responded to 1 µM norepinephrine with ΔF/F ≈ 0.2 (Fig S8B). The PF sensor is reported to have a *K*_D_ for cAMP of 7.2 μM (*58*), indicating that 1 µM norepinephrine elevated cAMP to the low μM range. Quantitative measurements in cultured cells reported an EC_50_ for PKA activation by cAMP of ∼5 μM (*47*). In our measurements PF responses (ΔF/F) were generally less than 50% of the saturating value, indicating cAMP concentrations < 7 μM, broadly within the physiological range.

Together, these experiments established the cAMP-SITES as useful tools for all-optical perturbation and readout of cAMP dynamics, provided that the bPAC concentration was maintained low enough to minimize perturbation to basal cAMP.

### All-optical measurements of PDE kinetics

We constructed a kinetic model for bPAC activation and the cAMP response (Fig. 2E) and derived expressions for the change in cAMP concentration as a function of time after a brief blue light pulse (**Supplemental Materials Section 3.1**).

We considered the response to a blue light pulse of duration short compared to the bPAC relaxation. After this pulse, the concentration of the bPAC activated state (bPAC*) followed an exponential decay with first-order rate constant *k*^bPAC^. We assumed that cAMP production is first order in bPAC*, and that substrate (ATP) is in excess. We further assumed that cAMP degradation is first order in cAMP and first order in PDE concentration, with rate *k*_PDE_ = *k*_cat_[PDE]/*K*_M_ (i.e. that the degradation is in the linear regime of the PDE Michaelis-Menten kinetics). PF responds to step-changes in cAMP concentration within ∼1 s (*58*), while the cAMP dynamics in our experiments lasted tens of seconds or longer, so we assumed local equilibrium of cAMP binding to PF. In these experiments we ensured PF ΔF/F < 1 to avoid saturating the PF sensor. Finally, we assumed that the PF does not substantially buffer the cAMP.

We transiently expressed all four cAMP-SITE constructs in separate dishes of HEK cells, delivered a wide-area brief blue light pulse (0.5 s, 10–21 µJ/mm^2^) to activate bPAC or bPAC^m^, and recorded the PF fluorescence responses (Fig. 2F). We fit the cell-average ΔF/F traces to our kinetic model (Methods) and found that for all four constructs, 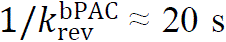, consistent with prior *in-vitro* measurements (*57*, *62*). We also found that 1/*k*_PDE_≈ 60–80 s for all combinations of membrane-bound and soluble bPAC and PF (Fig. 2G, Table 3). The kinetic fits almost perfectly overlaid the data for all four constructs (*R^2^* > 0.997). The consistency of *k*_PDE_ across variations in source and sensor sub-cellular localization suggests rapid equilibrium between membrane-proximal and intracellular cAMP, relative to the experimental timescale of 60 seconds. Since PF is not a ratiometric indicator, our experiments did not determine whether there was a constant multiplicative offset between intracellular and membrane-proximal pools; but the most parsimonious interpretation of the data was that the cAMP is well mixed throughout the cell.

### Cyclic AMP diffuses in the cytoplasm with *D* ≈ 130 µm^2^/s

Full mixing of cAMP across a ∼10 µm cell in ∼1 min could only occur if the cAMP diffusion coefficient in the cytoplasm obeyed *D* » 1 µm^2^/s. We thus sought to measure *D* for cAMP directly. We worked with MDCK cells because these cells often have a long and thin geometry, which allowed us to approximate cAMP transport as a 1-dimensional process. We expressed the cAMP-SITE construct PF^m^+bPAC^m^, selecting the membrane-bound bPAC to minimize diffusion of the cAMP source, and the membrane-bound PF to avoid dependence of the baseline fluorescence on cell thickness. Using the DMD, we transiently activated bPAC^m^ in a micron-scale region near one end of the cell, and then mapped the PF^m^ response (Fig. 3A–B, **Supplemental Video 1**). The PF^m^ fluorescence initially rose near the activated bPAC and then spread throughout the cell over ∼30 s.

**Figure 3.**
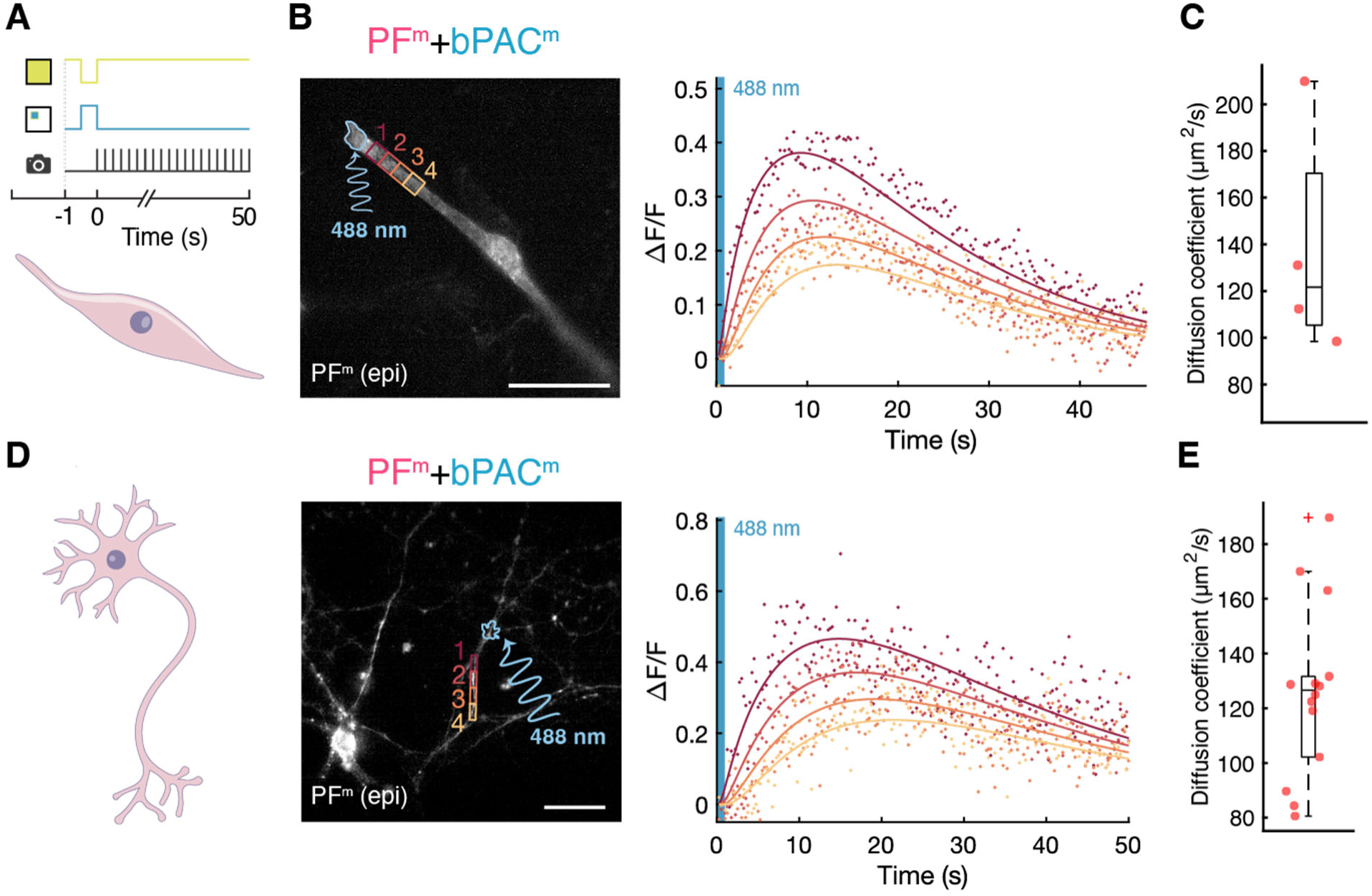
cAMP diffuses with *D* ∼ 130 μm^2^/s in MDCK cells and neurons. **A)** Illumination protocol for patterned bPAC^m^ activation (blue, 0.5 s) followed by wide-field PF^m^ imaging (yellow). **B)** Basal PF^m^ fluorescence in an MDCK cell expressing PF^m^ + bPAC^m^ taken with an epifluorescence microscope. 488 nm light was targeted to the indicated region (left). Dynamics of ΔF/F in the correspondingly colored regions were fit to the free diffusion model (*D* = 131 µm^2^/s, right). Scale bar: 50 µm. **C)** cAMP diffusion coefficient in MDCK cells: *D* = 134 ± 50 µm^2^/s (mean ± s.d., *n* = 4 measurements, 3 cells). **D)** Data from a representative cultured neuron (*D* = 81 µm^2^/s). Scale bar: 50 µm. **E)** cAMP diffusion coefficients in neurons: *D* = 126 ± 32 µm^2^/s, (mean ± s.d., *n* = 14 measurements, 9 cells). Boxes represent the interquartile range with the median denoted by the central line.

We augmented our kinetic model with a 1-D diffusion term for cAMP, leading to a four-parameter reaction-diffusion model which described the cAMP profile in time and space (**Supplemental Materials Section 4**). We performed a global fit of the model to the PF fluorescence dynamics over space and time within each cell, floating only the diffusion coefficient and a global scale factor (Fig. S9). We fixed the bPAC rate 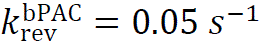, and fit ΔF/F for short times (*t* < 8 s), so the fit was not sensitive to *k*_PDE_ for 1/*k*_PDE_ » 8 s (recall, 1/*k*_PDE_ ≈ 60–80 s, Fig. 2G). The peak ΔF/F remained < 0.6, even in the regions closest to the bPAC activation, ensuring that the PF^m^ response did not saturate and that the cAMP concentration remained < 7 µM, based on the published dose-response curve of Pink Flamindo (*58*).

The best-fit cAMP diffusion coefficient was 134 ± 50 µm^2^/s, mean ± s.d., *n* = 4 measurements (Fig. 3C), almost identical to the results of Nikolaev *et al.* in cardiac myocytes (136 ± 36 μm^2^/s) (*25*). We performed the same measurement in dendrites of cultured rat hippocampal neurons (Fig. 3D; **Supplemental Videos 2, 3**) and found that the diffusion coefficient was similar, 126 ± 32 µm^2^/s, mean ± s.d., *n* = 14 measurements, 9 cells (Fig. 3E), consistent with prior experiments showing rapid agonist-evoked cAMP spread in cultured neurons (*64*). Measurements of free diffusion of other small molecules in cytoplasm yielded similar diffusion coefficients (*44*, *45*), implying that cAMP diffuses relatively freely in the cytoplasm of MDCK cells and neurons.

If there were nanoscale-localized membrane pools of cAMP, then the soluble (PF) and membrane-bound (PF^m^) reporters would be expected to show different diffusion coefficients. Matched experiments in dendrites expressing bPAC^m^ with either PF or PF^m^ as the cAMP sensor yielded similar diffusion coefficients; and in neither construct did the diffusion coefficient depend on ΔF/F_peak_ over the range 0.11 – 0.51 (i.e., the peak cAMP concentration; Fig. S10A–D). Together, these results are consistent with relatively free diffusion of cAMP, and with rapid equilibrium between any membrane vs. cytosolic pools.

As a separate test for small-scale compartmentalization, we searched for deviations in cAMP concentration around local clusters of bPAC. If individual bPAC molecules produced locally elevated concentrations of cAMP, then clusters would produce locally elevated patches of cAMP. In both HEK cells and neurons expressing GFP-bPAC^m^ we observed spatially heterogeneous GFP signals, as expected for a membrane-localized signal. We also observed some intracellular puncta, likely from bPAC^m^ molecules that had not yet trafficked to the membrane or had been internalized. However, upon a weak blue light stimulus, we observed no pixel-wise correlation between GFP-bPAC^m^ expression and PF^m^ ΔF/F in either HEK cells or neurons (Fig. S11, Pearson correlation coefficient *PCC* = 0.03 ± 0.18, *n* = 26 HEK cells; *PCC* = 0.10 ± 0.14, *n* = 8 neurons, mean ± s. d. Distribution medians were not significantly different from zero: *P* = 0.38, HEK cells, *P* = 0.11, neurons, Wilcoxon signed-rank test). In contrast, pixel-wise PF^m^ ΔF correlated strongly with baseline PF^m^ fluorescence, *F*, in both cell types (*P* = 9.3 × 10^-6^ for HEK cells; *P* = 0.0078 for neurons, Fig. S11C, F) confirming our ability to resolve fine-scale variations in PF^m^ fluorescence. We observed the same pattern for HEK cells expressing the soluble cAMP reporter PF and bPAC^m^ (Fig. S11). These results are, again, consistent with broadly distributed cAMP signaling, and inconsistent with sub-micron localization around cAMP sources.

### cAMP is geometrically compartmentalized in neurites

We next explored the interaction of cell geometry and cAMP reaction-diffusion dynamics. The AC sources of cAMP can be plasma membrane-bound or cytoplasmic (*65*). For example, GPCRs can signal either from the membrane or after becoming internalized (*66*, *67*). The PDE sinks of cAMP can also be plasma membrane-bound or cytoplasmic, and individual cells might have different isoforms of PDEs that are active at different cAMP concentrations (i.e. have different Michaelis constants *K*_M_) (*56*, *68–70*). Membrane-bound enzymes (AC or PDE) have a higher effective concentration in regions of higher surface-to-volume ratio, even if homogeneously distributed and homogeneously activated throughout the cell membrane. Hence the balance of AC and PDE activity could favor cAMP production in either soma or dendrites, depending on whether each enzyme is soluble or membrane-localized.

To test this idea experimentally, we compared the effects of bPAC^m^ vs. bPAC in cultured rat hippocampal neurons (Fig. 4A). In neurons expressing PF^m^+bPAC^m^, illumination with a wide-area pulse of blue light (0.5 s, 21–83 µW/mm^2^) led to larger peak cAMP concentrations in the dendrites than in the soma ((ΔF/F)_dendrite_/(ΔF/F)_soma_ = 1.6 ± 0.85, mean ± s.d., *n* = 62 dendrites, 6 neurons, Fig. 4B–D). In contrast, in neurons expressing soluble bPAC (PF^m^+bPAC), the same illumination and analysis protocol led to a larger PF^m^ signal in the soma than in the dendrites ((ΔF/F)_dendrite_/ (ΔF/F)_soma_ = 0.75 ± 0.22, mean ± s.d., *n* = 33 dendrites, 6 neurons, Fig 4E–G). The difference in the dendrite to soma ratio between cells expressing bPAC^m^ vs. bPAC was significant (*P* = 1.0e-8, Wilcoxon rank-sum test). Thus, membrane-bound bPAC preferentially produced cAMP in the dendrites, while soluble bPAC preferentially produced cAMP in the soma.

**Figure 4.**
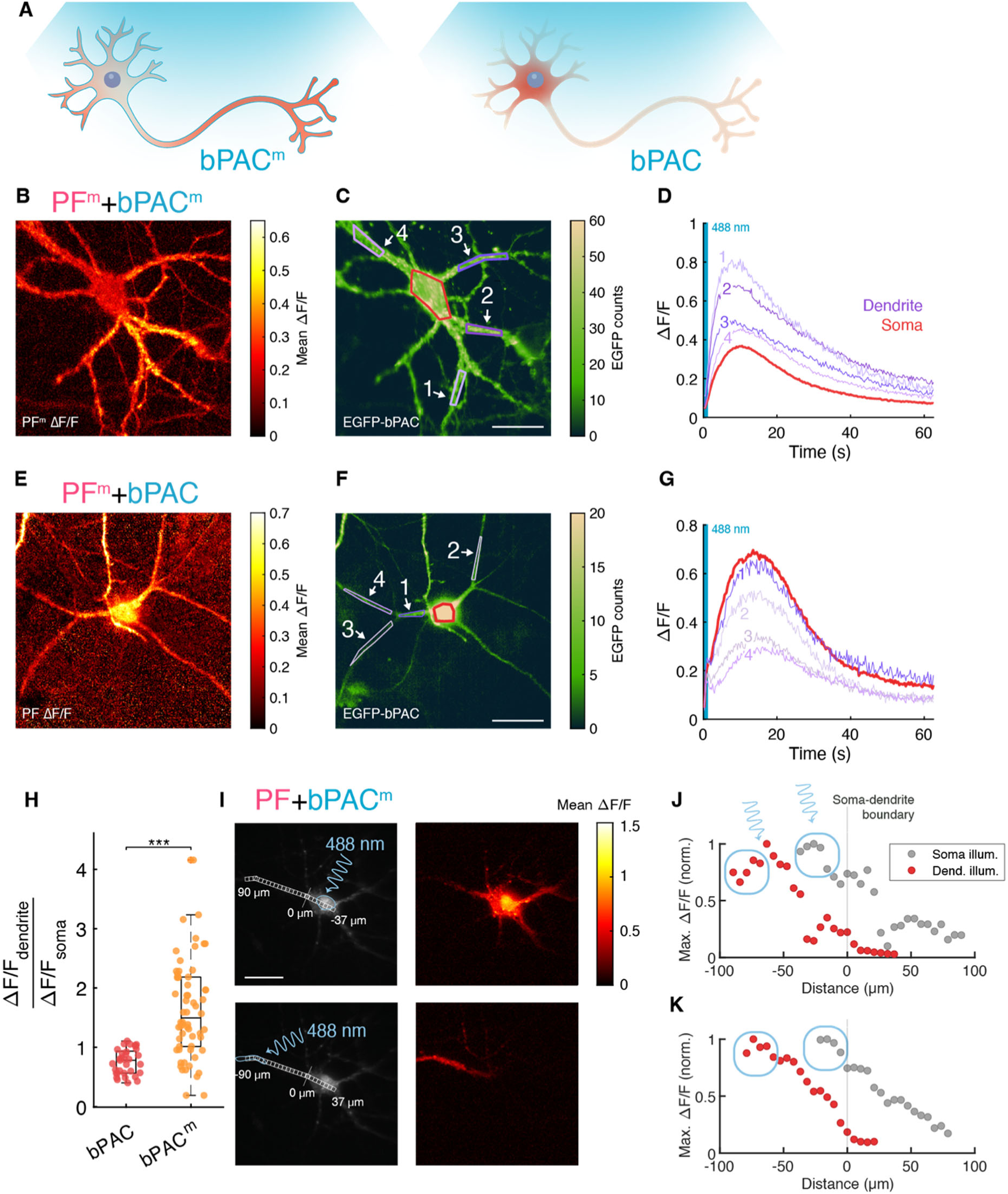
Membrane production and degradation cause heterogeneous cAMP distributions. **A)** Cartoon of a neuron expressing membrane-bound bPAC. Upon wide-area homogeneous blue light stimulation, cAMP accumulated preferentially in neurites due to a high surface-to-volume ratio and mixed bulk and membrane-localized degradation from PDEs. Soluble bPAC and the same illumination profile resulted in preferential cAMP accumulation in the soma due to its low surface-to-volume ratio and hence slower rate of cAMP degradation. **B)** A neuron expressing PF^m^+bPAC^m^ was exposed to a 0.5 s wide-area 488 nm light stimulus. Image shows mean PF^m^ ΔF/F during the 25 s following the stimulus. The cAMP increase in the neurites was larger than in the cell body. Scale bar: 50 µm. **C)** Epifluorescence GFP-bPAC^m^ image showing bPAC^m^ distribution. **D)** PF^m^ ΔF/F in the soma (red) and neurites (purple). **E–G)** Same as **B–D** but for a neuron expressing PF^m^+bPAC. **H)** Ratio of the peak ΔF/F in each dendrite scaled by the peak ΔF/F achieved in its corresponding soma (*n* = 62 dendrites and 6 cells for bPAC^m^, and *n* = 33 dendrites and 6 cells for bPAC. *** *P* = 1.0 × 10^-8^, Wilcoxon rank-sum test). Boxes represent the interquartile range with the median denoted by the central line. **I)** Neurons expressing PF+bPAC^m^ were subject to 0.5 s of 488 nm patterned illumination in the region circled (blue, either soma or dendrite). Right: Mean PF ΔF/F for the first 25 s after stimulus. Scale bar: 50 µm. **J)** Maximum PF ΔF/F achieved in each region along the line denoted in (**I**), normalized by the peak value. For each stimulus location, distance was measured relative to the blue illumination spot, and offset so zero corresponds to the boundary between the soma and dendrite. Soma-produced cAMP permeated the dendrites, but dendrite-produced cAMP did not change cAMP concentration in the soma. Blue circles indicate illumination spots. Data are discretized into in 5.3 µm segments. **K)** Same as **(J)**, but for another soma-dendrite pair in a different neuron.

We then studied the interchange of cAMP between dendrites and the soma. In neurons expressing PF+bPAC^m^, we used patterned blue light (0.5 s, 83 µJ/mm^2^) to activate cAMP production in a dendrite, and we mapped the distribution of cAMP along the line spanning from the dendrite to the middle of the soma (Fig. 4I). The change in cAMP concentration declined smoothly to approximately zero at the junction of the dendrite and the soma, a consequence of the comparatively large volume of the soma causing dilution of cAMP sourced from the dendrite. Essentially, the soma acted as an absorbing boundary at the junction with the dendrite, confirming a previous theoretical prediction (*36*).

We then patterned the blue light to activate cAMP production in the soma of the same cell. In this case, the cAMP propagated ∼20 μm into the dendrites. The soma was able to provide cAMP to the dendrites with minimal dilution of the source (Fig. 4J, K). Hence cAMP transport is asymmetric with the soma influencing the proximal dendrites more than the proximal dendrites influenced the soma, a simple consequence of the different volumes of these structures. For distal dendrites, the increased volume in regions toward the soma as well as the branching structure of the dendritic tree could further compartmentalize cAMP signals.

### The balance of diffusion and degradation sets a cAMP signaling length

Finally, we measured the characteristic length-scale for cAMP transport (the Thiele length) in neurites, set by the balance of cAMP diffusion and PDE-mediated degradation (Fig. 5A). In cultured neurons expressing either PF^m^+bPAC^m^ or PF+bPAC^m^, we repeatedly targeted the blue light to a small patch of neurite and mapped the cAMP concentration until it reached steady state (Fig. 5B–D). We fit the steady state concentration profiles to an exponential decay vs. position (Fig. 5E).

**Figure 5.**
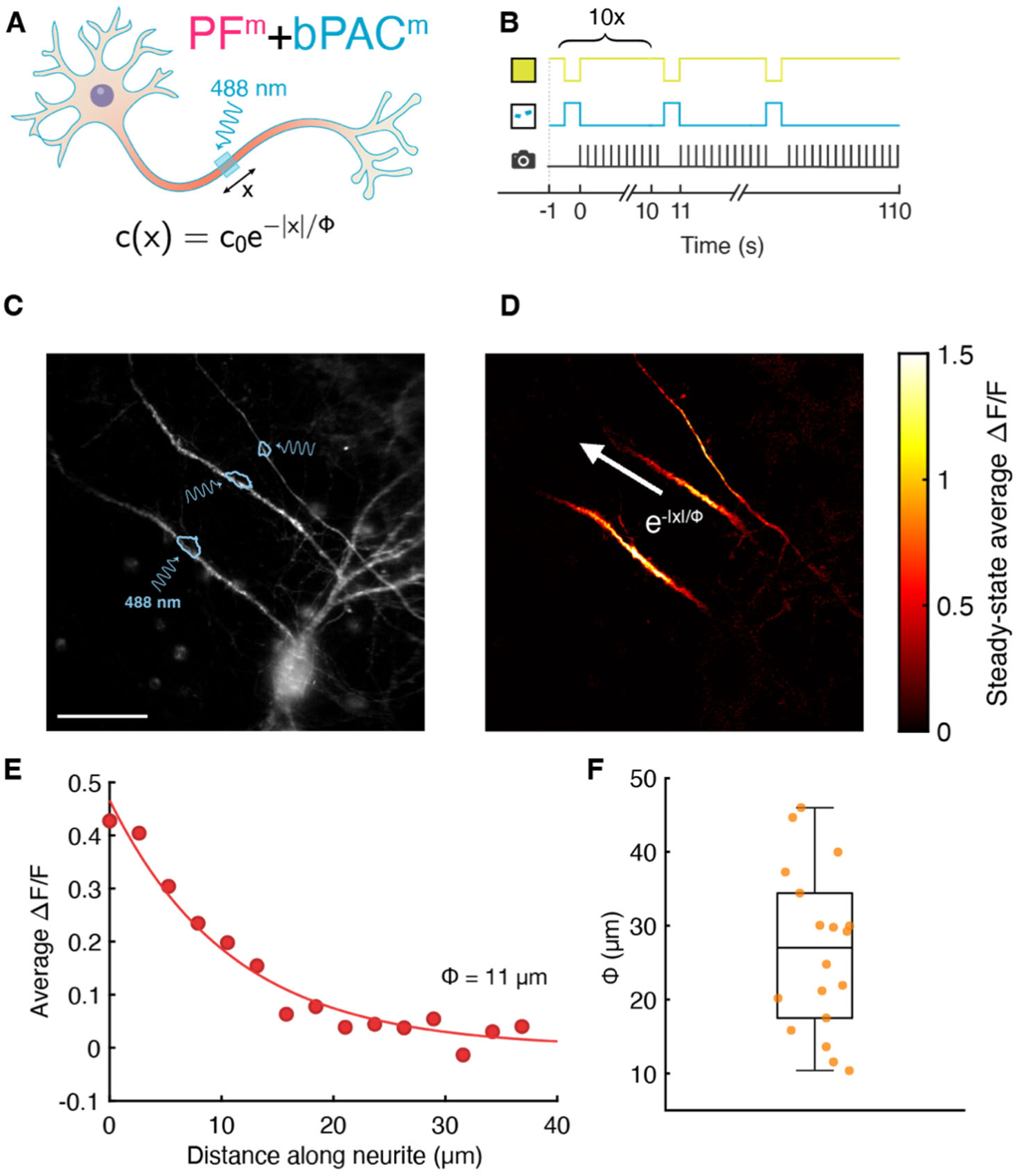
cAMP length constants in neurites. **A)** Cartoon of localized cAMP production in a neurite. The steady-state cAMP concentration profile decays exponentially with Thiele length, ϕ, set by the balance between diffusion and degradation. **B)** Illumination protocol comprising interlaced localized blue light stimulation and wide-field yellow light imaging. **C)** PF^m^+bPAC^m^-expressing neurons were subject to targeted bPAC stimulation and cAMP profile was imaged with an epifluorescence microscope at steady state. Image is of basal PF fluorescence. Scale bar: 50 µm. **D)** Mean PF ΔF/F after steady state was achieved. **E)** Mean ΔF/F along the neurite, with the illumination region at position 0. Space is discretized in 2.6 µm segments. An exponential fit yielded 1/ϕ = (0.091 ± 0.015) µm^-1^. **F)** Thiele length ϕ = 27 ± 11 µm (*n =* 18 measurements, 13 cells, 4 dishes, mean ± s. d.).

We reasoned that if membrane-tethered PDEs created localized nanodomains that did not interconvert with the cytoplasmic pool, then a membrane-tethered reporter (PF^m^) would exhibit significantly different spatial profiles compared to a soluble reporter (PF). The Thiele length was ϕ = 27 ± 11 µm for the membrane-bound reporter (mean ± s.d., *n* = 18 measurements; Fig. 5F) and ϕ = 24 ± 9 µm for the soluble reporter (*n* = 11 measurements, Fig. S10E–G). The similar Thiele lengths for soluble vs. membrane-bound cAMP reporters is, again, consistent with rapid equilibrium between these pools.

The Thiele length was independent of peak cAMP concentration over the range of (ΔF/F)_peak_ from 0.19 – 0.48 (Fig. S10G, H), indicating that nonlinear feedback mechanisms (cAMP buffer saturation, cAMP-dependent PDE activation) were not modulated in these experiments. The much greater length-scale of cAMP signaling (∼25 μm) compared to the dendrite diameter (< 6 μm) supports the interpretation that there are no meaningful cAMP gradients transverse to the dendrite axis, so the diffusion can be modeled as 1-dimensional.

The slow inactivation rate of activated bPAC 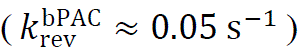 prevented direct measurement of the (much faster) rate of PDE-mediated cAMP recovery in neurons. However, from the diffusion coefficient (*D* ≈ 126 μm^2^/s), and length-scale (ϕ ≈ 27 µm) we estimate 1/*k*_PDE_ = ф^2^/*D* ≈ 5.8 s. This timescale is broadly consistent with previously reported cAMP recovery times in mammalian dendrites (*71*, *72*). Thiele length measurements provide a convenient way to assess timescales of PDE action from steady-state measurements.

### Scaling relations for small-molecule transport in cells

Detailed numerical models have been constructed to simulate cAMP transport in neurons (*38*), myocytes (*39*), and endothelial cells (*40*); and sophisticated analytical approaches have also been applied to estimate cAMP transport in various geometries (*21*, *36*). While thinking about cAMP transport in dendrites, we found it useful to have simple scaling relations as a guide to intuition. Similar ideas have been explored theoretically and experimentally with other diffusible messengers (*73–75*), though we are not aware of a prior unified presentation of the scaling relations shown in Fig. 6.

**Figure 6.**
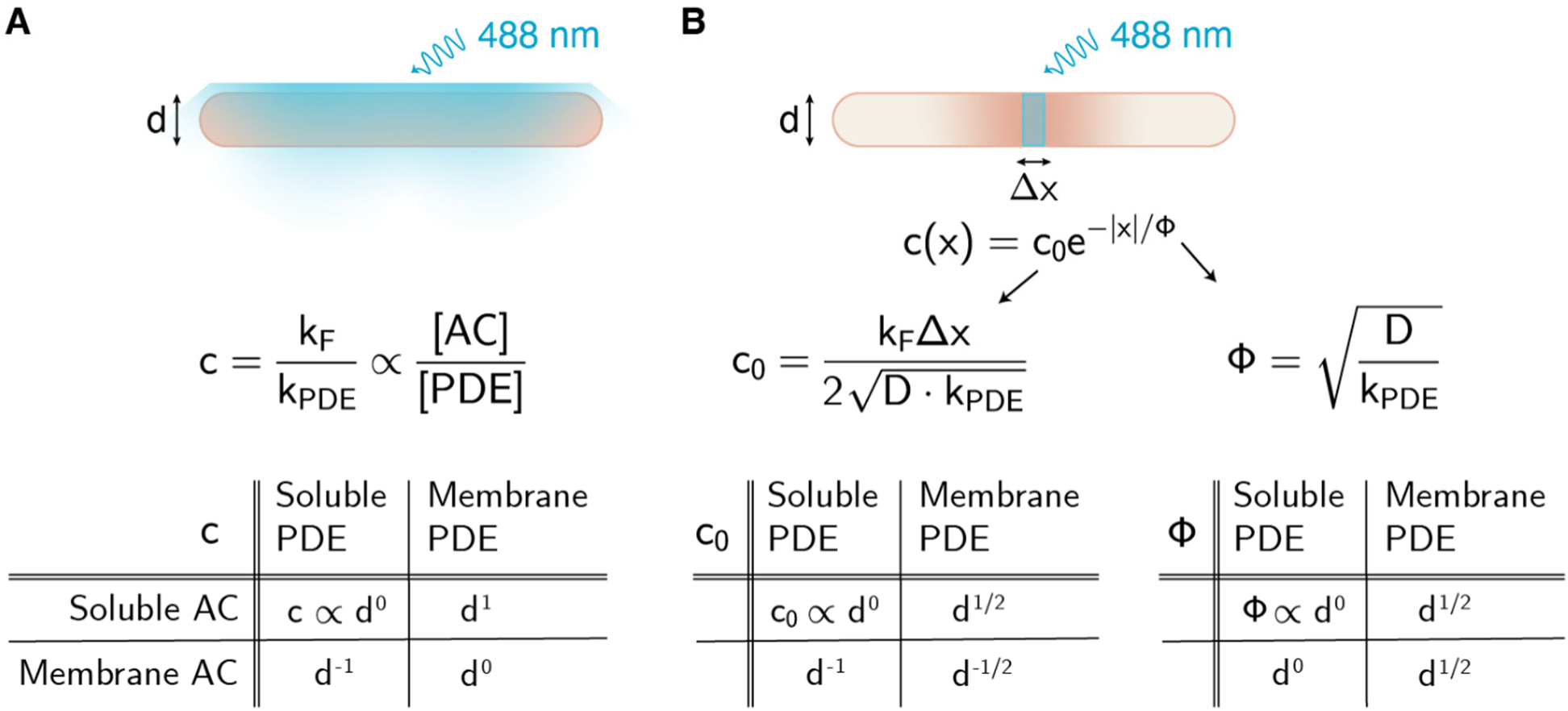
Scaling of cAMP concentration and diffusion length with tube diameter and enzyme localization. **A)** Cartoon of wide-area homogeneous bPAC activation in a tube with diameter *d*. The steady state cAMP concentration depends on the ratio between AC and PDE effective concentrations. Effective concentrations of membrane-bound AC or PDE scale as *d*^-1^ (assuming constant surface density); effective concentrations of soluble enzymes scale as *d*^0^. **B)** Cartoon showing localized bPAC activation in a region with small width Δ*x*. The concentration follows a decaying exponential in space, and both the peak concentration *c*_0_ and the length-scale ф can vary with tube diameter depending on enzyme localization.

We modeled cAMP dynamics in a tube-shaped process as a one-dimensional reaction– diffusion system with a first-order degradation rate. The full model and calculations are in the **Supplemental Materials, Sections 5–6.** The key geometrical observation is that the effective concentration of a membrane-bound enzyme (AC or PDE) is proportional to the surface-to-volume ratio, i.e. is higher for thinner tubes. Consequently, even when enzyme activities are spatially uniform, membrane localization can bias steady-state cAMP toward thinner or thicker processes depending on whether production (AC) and degradation (PDE) are each predominantly soluble or membrane-localized (Fig. 6A).

Membrane localization of can also affect the length-scale of cAMP spread from a spatially localized source (the Thiele length, ф). The key result is that ф depends on the localization of the sink: for predominantly soluble PDE activity, ф is predicted to be approximately independent of tube diameter, whereas for predominantly membrane-bound PDE activity, ф scales as 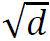, where *d* is the tube diameter (Fig. 6B). The localization of the source does not affect ф.

These scaling relations are qualitatively consistent with our data comparing cAMP profiles evoked by bPAC^m^ (which favored the dendrites) vs. bPAC (which favored the soma; Fig. 4). Similarly, Bacskai and coworkers showed that bath-applied serotonin preferentially elevated cAMP in *Aplysia* dendrites relative to soma, a consequence of membrane-localized AC activation (*22*). In hippocampal neurons, some pharmacological perturbations preferentially elevate cAMP in the distal dendrites (*14*, *38*, *72*), while others act primarily on the soma (*72*), presumably because of differential effects on membrane vs. internal cAMP-processing enzymes.

In our experiments, the Thiele length did not show clear correlation with diameter for neurite diameters ∼0.5 – 6 µm (Fig. S12). This observation is consistent with a primarily soluble pool of PDEs, but is also consistent with a dominant influence of factors not included in our simple model. Possible contributors to variation in effective Thiele length include variations in neurite geometry (e.g. dendritic spines can decrease effective axial diffusion (*76*, *77*)), and cell-to-cell or sub-cellular variations in PDE expression level, activity, or localization. A direction for future research will be to determine the mechanisms driving deviations from simple scaling in sub-cellular cAMP transport.

### Spatial logic of chemical vs electrical signaling

The reaction-diffusion equation governing cAMP transport has same mathematical structure as the cable equation for propagation of membrane voltage, an analogy previously noted by others (*42*, *50*, *78*). However, the dependence of the parameters on tube diameter can be different. In the electrical case, current sources and sinks are always across the membrane, and so the scaling rules are the same as for the case where AC and PDE are both membrane-bound. The electrical length constant 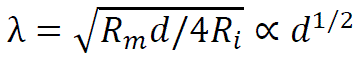, where *R_m_* is the membrane resistivity and *R_i_* is the axial resistivity. Typically electrical length constants in dendrites are λ > 150 μm (*79*, *80*), longer than most chemical signaling lengths. However, if the PDE is soluble, then φ ∝ *d*^0^and has different scaling with *d* compared to *λ* ∝ *d*^1/2^. Then there is, in principle, a crossover point where chemical signaling becomes longer-range than electrical signaling. If we posit a 6-fold difference between λ (∼150 μm) and ϕ (∼25 μm) at *d* = 5 μm, then a 36-fold decrease in *d* would be required to achieve parity (assuming purely soluble PDE activity). A diameter of 140 nm is attainable in cilia and cytoskeleton-free membrane tubes (*81*), but otherwise is outside the physiologically relevant range.

## Discussion

Some have argued that cAMP diffuses relatively freely in cells (*25*, *82*, *83*), while others have suggested that highly restricted or buffered diffusion generates nanoscale compartments (*30*, *35*), and that individual GPCRs with closely coupled downstream effectors can function independently of other GPCRs (*32*). Using patterned cAMP production, we observed transport of cAMP from a spatially localized source and made quantitative measurements of its diffusion and transport lengths. We compared membrane-bound and soluble combinations of actuator and reporter and found little difference between the constructs in terms of cAMP degradation kinetics or transport within dendrites. Fine-grained measurements of the relation between bPAC distribution and PF response showed no evidence of nanoscale cAMP compartmentalization, consistent with the ∼25 µm signaling length of cAMP.

A potential criticism of this work is that the cAMP concentrations probed (we estimate 1 – 10 μM) may be higher than found in neurons, and thus localization mechanisms active at lower concentrations may have been overwhelmed. We think this argument is unlikely for several reasons. (1) Buffering does not decrease localization lengths, because buffers equally suppress diffusion and degradation. (2) We did not observe strong concentration-dependent trends in *D* or ϕ, even at low ΔF/F, suggesting that extrapolation to lower cAMP concentrations would yield similar results. (3) While basal cAMP concentrations can be in the tens of nM, peak concentrations frequently reach several μM, within the range of our experiments (*38*, *72*). Furthermore, recent studies measured physiological cAMP signals in mouse hypothalamic and parabrachial neurons (*15*, *63*) using the cAMP sensor cADDis, which is less sensitive than Pink Flamindo (*K_D_* > 10 µM (*84*) versus *K_D_* = 7.2 µM). We propose that cAMP differential signaling is primarily governed by membrane-delimited compartmentalization in sub-cellular domains, combined with differential scaling of soluble vs. membrane-bound enzyme activities as a function of local surface-to-volume ratio.

The length-scale of cAMP transport in neurons (∼25 μm) was short enough that cAMP remained compartmentalized in individual dendrites. Therefore, it is plausible that individual portions of the dendritic tree could experience localized neuromodulatory effects, possibly mediating local changes in dendritic computations or plasticity rules. The length-scale of cAMP transport is in between that of Ca^2+^ (several microns) (*42*, *50*, *85*) and membrane voltage (typically ∼150 μm) (*79*, *80*). Further, our results show that drugs that differentially target membrane-bound versus cytoplasmic PDE (or AC) populations have potential to modulate the size and concentration of cAMP domains and therefore to have differential effects on downstream signaling.

Direct AC stimulation, as in cAMP-SITES, acts downstream of GPCRs. The precise spatial map of AC activity after a physiological GPCR stimulus is unknown, though the timescale is set by diffusive search between a Gα_s_ subunit and an AC molecule (*86*). Our measurements of diffusion and length scale should be interpreted as impulse responses to a defined and restricted AC stimulus, which can then be convolved with various upstream stimulus patterns.

The cAMP-SITES tools we developed here could be broadly useful for studying sub-cellular and intercellular cAMP transport. For example, the bPAC and the Pink Flamindo could be independently localized to different regions, such as mitochondria, nucleus, dendritic spines or flagella; one could thereby probe cAMP transport between any two of these regions. The bPAC and Pink Flamindo could also be expressed in adjacent cells, permitting studies of cAMP transport through gap junctions. The cAMP-SITES provide a convenient and robust assay of PDE activity, which may have use in high-throughput drug or genetic screens (*87*).

The effects of surface-to-volume ratio discussed here apply broadly to other small molecules and to systems where interacting partners are localized differentially on the membrane versus in the bulk (*75*, *88*, *89*). An important goal for future research will be to characterize the diffusion coefficients, length constants, and time constants for other signaling molecules. There exists a large and rapidly growing toolkit of spectrally distinct optogenetic perturbation tools and fluorescent reporters (*90–95*), and finding compatible pairs will open new windows into the spatial aspects of sub-cellular signaling.

## Supporting information

Supplementary modeling and calculations

## Acknowledgments

We thank Andrew Preecha, Camila Bodden, and Shahinoor Begum for technical assistance. This work was supported by NSF Quantum Sensing for Biophysics and Bioengineering (QuBBe) Quantum leap challenge institute (QLCI) grant OMA-2121044, a Hertz Foundation Fellowship (KX), the NSF Graduate Research Fellowship Program under Grant #2140743 and #1745303 (KX), and a Vannevar Bush Faculty Fellowship N00014-18-1-2859 (AEC).

## Declaration of Interests

AEC is a founder of Luminos LLC. The other authors declare no competing interests.

## Supplemental Materials

Quantitative modeling of cAMP kinetics and diffusion.

## Supplemental Video Captions

**Video S1: cAMP diffusion in MDCK cells.** Video shows membrane-bound Pink Flamindo (PF^m^) fluorescence response, ΔF, to focal blue light activation of membrane-bound bPAC (bPAC^m^). The PF^m^ response is overlaid on a greyscale image of the MDCK cell.

**Videos S2, S3: cAMP diffusion in neuronal dendrites.** Video shows membrane-bound Pink Flamindo (PF^m^) fluorescence response, ΔF, to focal blue light activation of membrane-bound bPAC (bPAC^m^). The PF^m^ response is overlaid on a greyscale image of the neuronal dendrites.

## Materials and Methods

### Genetic constructs

The genes were cloned into a second-generation lentiviral backbone with cytomegalovirus (CMV) promoter using standard Gibson Assembly. Briefly, the vector was linearized by double digestion using restriction enzymes (New England Biolabs, Ipswich, MA) and purified by the GeneJET gel extraction kit (ThermoFisher, Waltham, MA). DNA fragments were generated by PCR amplification and then fused with the backbones using NEBuilder HiFi DNA assembly kit (New England Biolabs). Constructs were transformed and amplified in NEB 5-alpha Competent E. coli (New England Biolabs), and isolated using GeneJET miniprep kit (Thermo Scientific, Waltham, MA). All plasmids were verified by sequencing (GeneWiz, Cambridge, MA). The plasmids used in this work are listed in Table 2 and are available at Addgene.

### Cell culture and expression

Human embryonic kidney (HEK293T) cells (ATCC CRL-11268) were cultured and transfected following standard protocols. Briefly, HEK293T cells were grown at 37 °C, 5% CO_2_, in Dulbecco’s modified Eagle’s medium (DMEM) supplemented with 10% fetal bovine serum (FBS), penicillin (100 U/mL), and streptomycin (100 µg/mL). For maintaining or expanding the cell culture, we used tissue culture (TC)-treated culture dishes for optimum growth of anchorage-dependent cells (Corning). For imaging, cells were plated on poly-D-lysine (PDL)-coated glass-bottomed dishes (Cellvis, Cat.# D35-14-1.5-N). 2 µg of plasmid DNA was transfected per dish using Transit 293T (Mirus) following the manufacturer’s instructions and measured at least 48 h later.

Madin-Darby canine kidney (MDCK) cells were cultured at 37 °C, 5% CO_2_, in DMEM supplemented with 10% FBS, penicillin (100 U/mL), and streptomycin (100 µg/mL). We used the TC-treated culture dishes (Corning) for maintaining or expanding the cell culture, and uncoated glass-bottomed dishes for imaging (Cellvis, Cat.# D35-14-1.5-N). MDCK cells were transduced with 200 µL lentivirus.

Primary E18 rat hippocampal neurons (fresh, never frozen, BrainBits #SDEHP) were dissociated following vendor protocols and plated in PDL and laminin-coated glass bottom dishes. Neurons (21k/cm^2^) were cocultured with primary rat glia (27k/cm^2^) to improve cell health and maturation. Neurons were transduced after 5-10 days in culture with 200 µL of low-titer lentivirus-containing media. Neurons that were co-transfected were transduced after 5 days in culture with 25 µL of lentivirus containing GFP-bPAC and 75 µL of lentivirus containing PF, and imaged 3–4 days after.

### Lentivirus production

Lentivirus preparations were made in house. HEK293T cells were grown to 80% confluence in a 35 mm dish. Fresh DMEM was exchanged 1 to 2 hours before DNA transfection. Transfer plasmid (0.9 µg), second generation packaging plasmid psPAX2 (0.9 µg; Addgene #12260), and envelope plasmid VSV-G (0.6 µg; Addgene #12259) were mixed with reduced-serum Minimal Essential Medium (Opti-MEM; 250 µL). The mixture was vortexed briefly and incubated at room temperature for 20 min. The mixture was then added dropwise to the dish of HEK293T cells and incubated for 48 hours. Cell culture supernatant was collected and filtered using a 0.45-μm syringe filter and aliquoted for storage at −80°C.

### Imaging

Before optical stimulation and imaging, cultured cells in 35 mm dishes were washed twice with 1 mL extracellular (XC) buffer containing (in mM): 125 NaCl, 2.5 KCl, 2 CaCl_2_, 1 MgCl_2_, 15 HEPES, 25 sucrose (adjusted to pH 7.3 and 304 mOsm L^-1^) and then incubated with 1 mL XC buffer warmed to 37 °C. Samples were imaged at room temperature.

Experiments were conducted on a home-built inverted fluorescence microscope equipped with 405 nm, 488 nm, 532 nm, 561 nm, 594 nm, and 640 nm laser lines and a scientific complementary metal-oxide semiconductor (sCMOS) camera (Hamamatsu ORCA-Flash 4.0). Beams from lasers were combined using dichroic mirrors and sent through an acousto-optic tunable filter (AOTF; Gooch and Housego TF525-250-6-3-GH18A) for temporal modulation of intensity of each wavelength. The beams were then expanded and sent either to a DMD (Vialux, V-7000 UV, 9515) for spatial modulation or sent directly into the microscope (to avoid power losses associated with the DMD). The beams were focused onto the back-focal plane of a 60×/1.2-NA (numerical aperture) water-immersion objective (Olympus UIS2 UPlanSApo ×60/1.20W) or a 20×/0.75-NA objective (Olympus UIS2 UPlanSApo ×20/0.75). Fluorescence emission was separated from laser excitation using a dichroic mirror (488/561/633). Imaging of Pink Flamindo fluorescence was performed with a 561 nm laser at an illumination intensity of 7.4 mW mm^−2^ and imaged with a 596/40 emission filter. Stimulation of bPAC was performed with 488 nm laser at illumination intensities of 19–283 µW mm^−2^. Photoactivation of paGFP was performed with 405 nm laser for 10 seconds. Imaging of GFP fluorescence was a terminal experiment (due to concurrent bPAC activation) with 488 nm laser at an illumination intensity of 21 µW mm^−2^ and a 532/50 emission filter.

Confocal imaging of fixed cells was performed on a Zeiss LSM 980 microscope. Cells were fixed with 5% paraformaldehyde and shielded from light until imaging. Confocal imaging of live cells was performed on a Zeiss LSM 900 microscope.

### Modeling and analysis

Data were analyzed in MATLAB. All videos were collected at 4 frames/s. ΔF/F was computed by taking the cell-averaged raw fluorescence intensities after subtracting the camera offset and background counts, then dividing by the time-averaged baseline fluorescence. For diffusion and Thiele length analyses, the background was computed by taking the 35^th^ percentile of fluorescence intensity for pixels in a dark region of the field of view during imaging. For kinetic calculations, the background was computed by taking the mode of all pixels during the baseline period after subtracting camera dark counts. For videos and time-averaged images of the entire field of view, the background was not subtracted for visualization purposes. Time-averaged neurite length-scale images such as in Fig. 5 were 2D median filtered with a kernel size of 3×3 pixels. For neuronal forskolin responses in Fig. S8, cells were only included if their pixel-wise response (ΔF vs. F) to forskolin stimulation was linear, with *R*^2^ *>* 0.1

Kinetics, diffusion, and neuron length-scale modeling as well as further information about analysis can be found in the **Supplemental Materials**.

## Supplemental Figures

**Figure S1.**
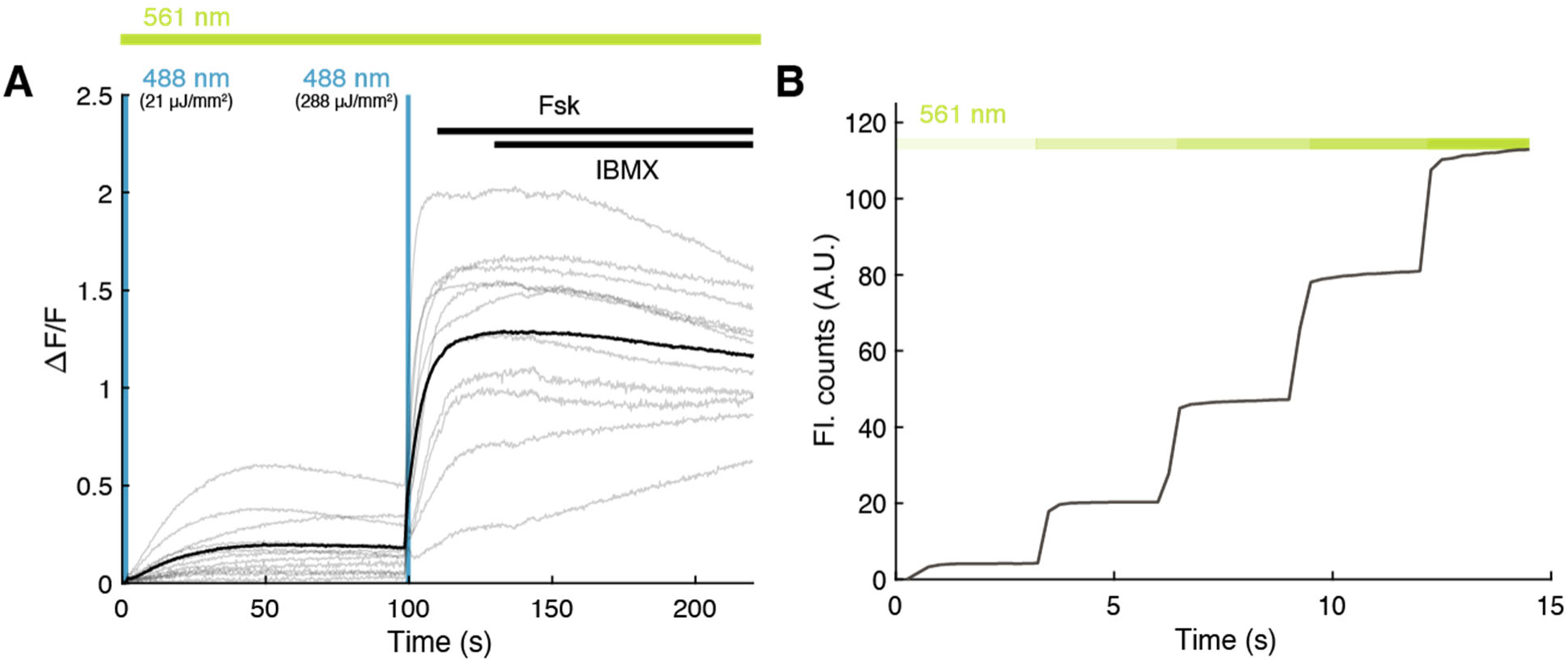
Characterization of PF+bPAC cAMP-SITE. **A)** Strong bPAC activation saturated the PF reporter. In HEK cells expressing bPAC+PF, a low-intensity 0.5 s pulse of 488 nm light activated bPAC and evoked a PF response. At 100 s, a high-intensity 0.5 s pulse of 488 nm light evoked a larger PF response. After 10 s we added forskolin (final concentration 100 µM), then IBMX (final concentration 200 µM). These drugs did not further increase the PF response, implying saturation of the PF reporter. **B)** Yellow light did not activate bPAC. HEK cells expressing bPAC+PF were exposed to step-wise increases of 561 nm light, from 2 mW/mm^2^ to 10 mW/mm^2^. The PF fluorescence showed step-wise increases proportional to illumination intensity. Absence of slow PF transient responses confirmed negligible changes in cAMP concentration during this protocol. Plot shows the average of *n* = 15 cells.

**Figure S2.**
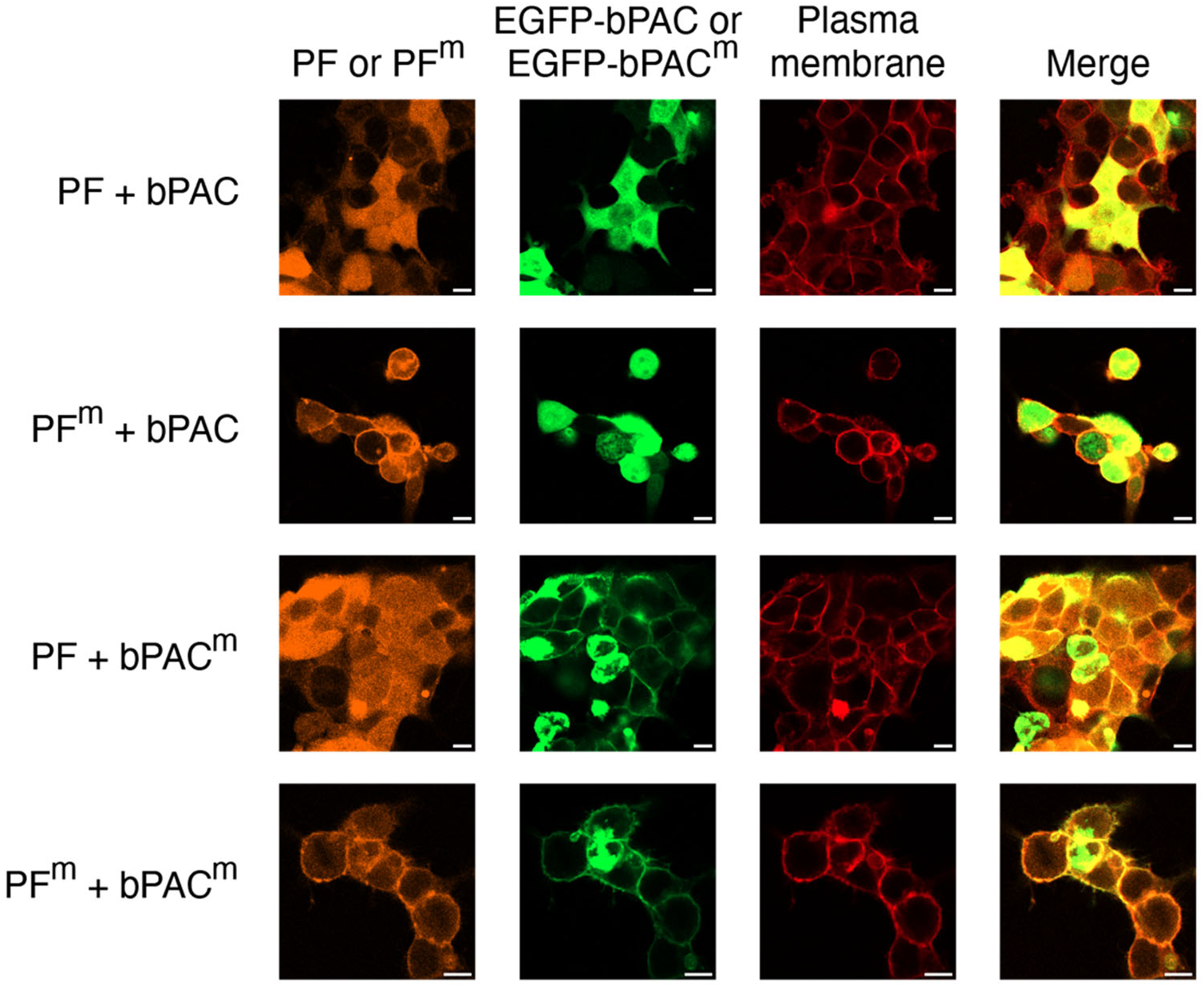
Localization of all four cAMP-site constructs in HEK293T cells. Localization of PF (soluble or membrane bound via myristoylation), GFP-bPAC (soluble or membrane bound via the CAAX domain), Cellmask Deep Red to label the plasma membrane, and the merged image. The first two columns for PF+bPAC and PF^m^+bPAC^m^ are the same as in Fig. 2. Scale bars 10 µm. All images are single-plane confocal images of cells transfected with p2a constructs.

**Figure S3.**
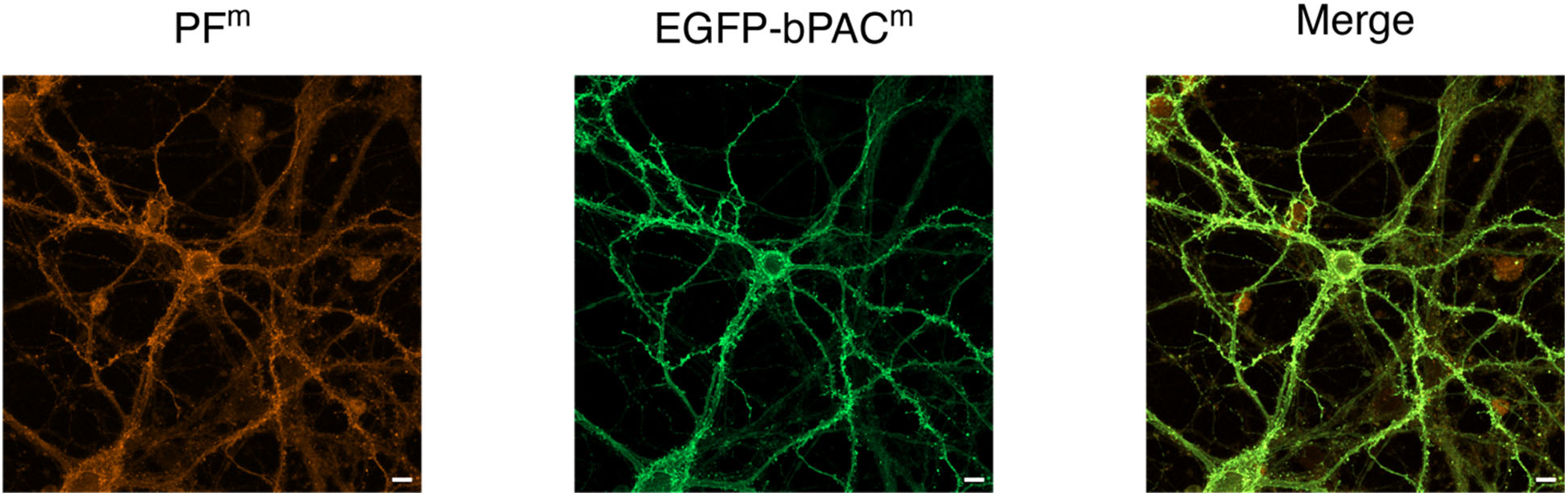
Membrane localization of Pink Flamindo and bPAC in neurons. Confocal images of localization of membrane-bound PF and membrane-bound bPAC in neurons. Images are orthogonal projections which include the apical and basal membrane. Neurons were transfected with p2a lentiviral constructs. Scale bar: 10 µm.

**Figure S4.**
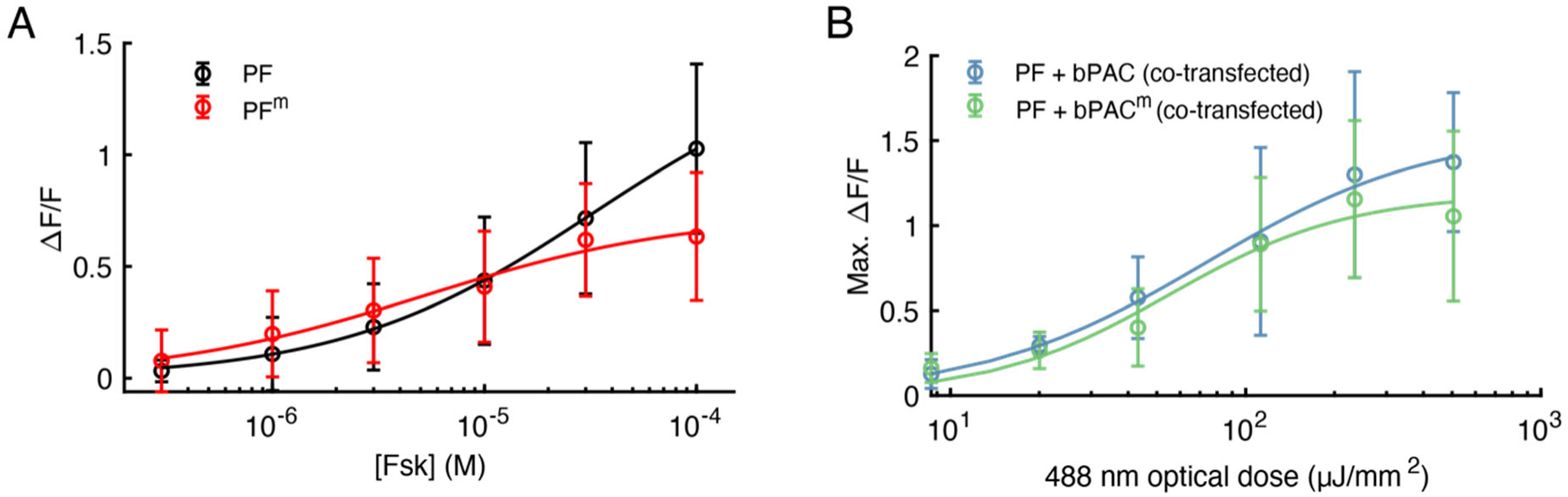
Validation of soluble and membrane-targeted Pink Flamindo and bPAC. **A)** Dose-response curve of soluble and membrane-tethered Pink Flamindo to forskolin in HEK cells. Data represent mean ± s.d. (*n* = 65 cells for PF and 37 cells for PF^m^, pooled from two experiments). **B)** Dose-response curve of PF ΔF/F to blue light, for cells expressing both PF and either bPAC or bPAC^m^. Cells included were between the 5^th^ and 50^th^ percentiles of bPAC or bPAC^m^ expression and had basal PF fluorescence < 100 counts. Blue light doses ranged from 8.6 – 506 µJ/mm^2^. Each point represents an average of single cell responses. Each field of view was exposed to blue light only once. Data represent mean ± s.d. (*n* > 50 cells for each point, from 2 experiments).

**Figure S5.**
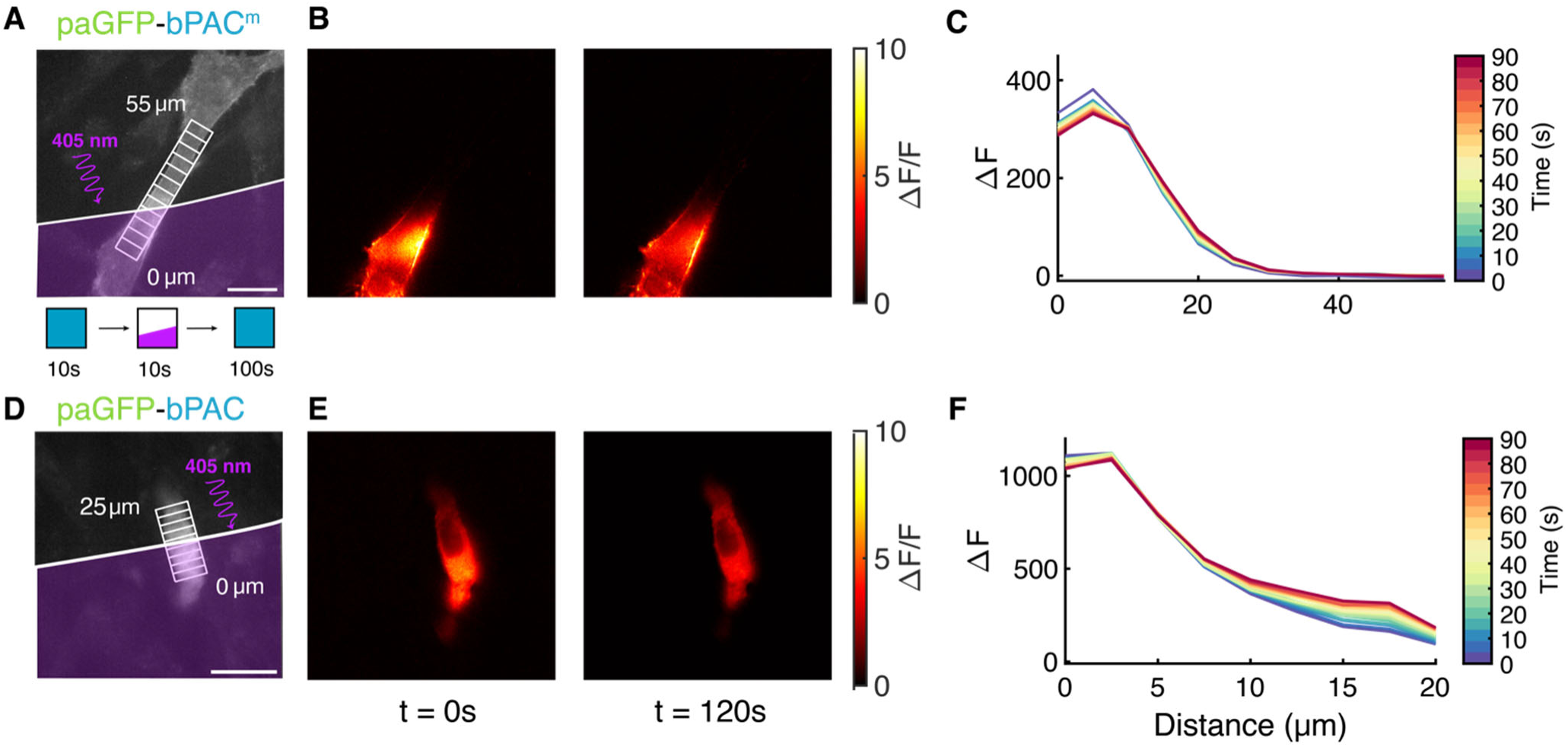
Photoactivation probes diffusion of membrane-bound and soluble bPAC. **A)** paGFP tagged to bPAC^m^ was activated with purple (405 nm) light in a subcellular region for 10 s. GFP fluorescence was recorded with blue (488 nm) light. Scale bar: 20 µm. **B)** Images of GFP ΔF/F. Left: at t = 0 (immediately after the end of stimulation with purple light). Right: t = 120 s. **C)** ΔF for each region in **A** color-coded by time from t = 0 to t = 90 s. **D–F)** Same as **A–C** but for soluble bPAC. For paGFP-bPAC^m^, after 25 s only 5% of the fluorescence (quantified by ΔF) had spread outside the photoactivated region. For the soluble paGFP-bPAC, 25% of the fluorescence had spread during the same time interval (*n* = 3 cells for each construct).

**Figure S6.**
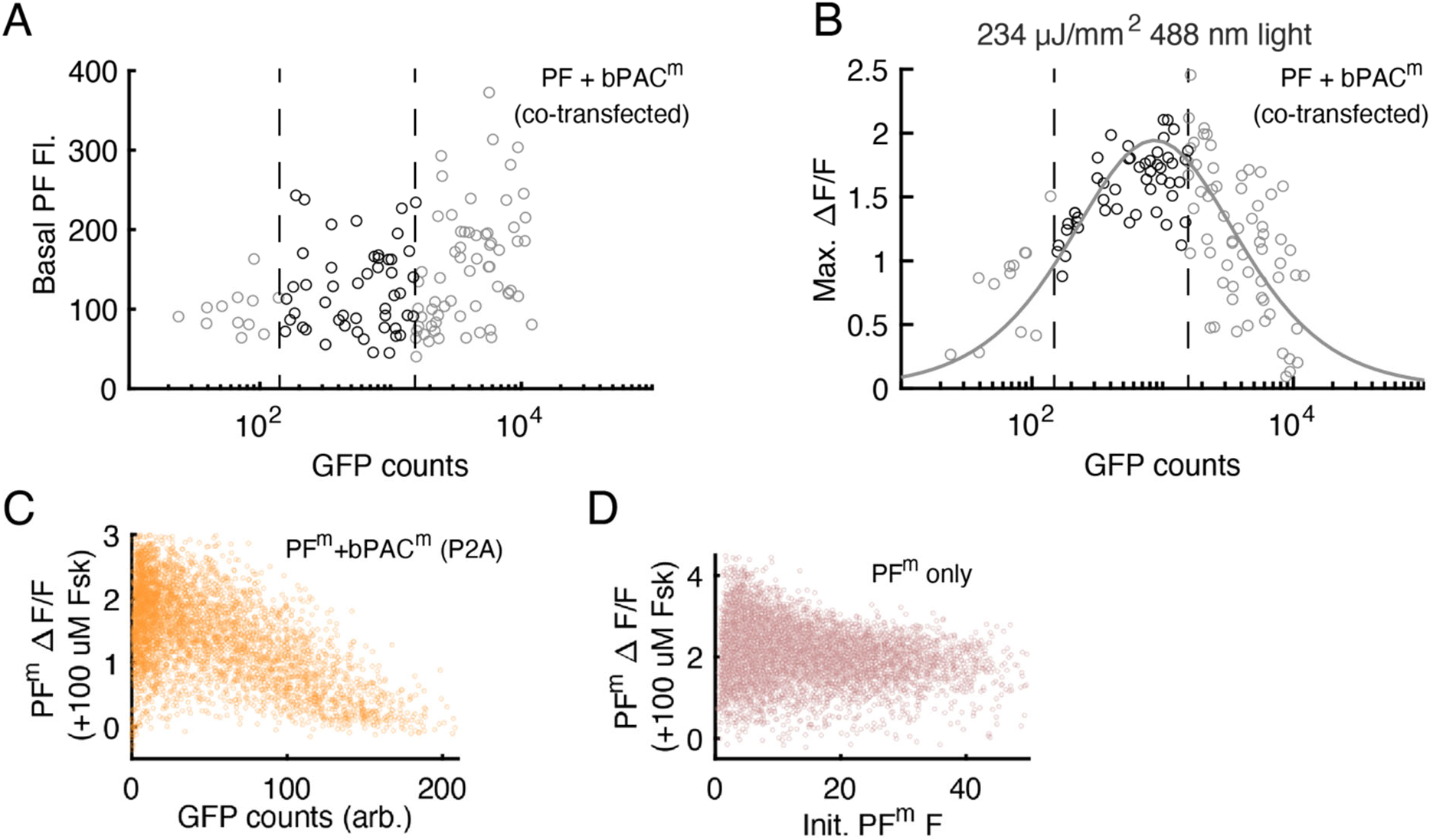
bPAC and bPAC^m^ dark-state activity. **A)** Cell averaged basal PF fluorescence vs. GFP fluorescence for HEK cells expressing PF and GFP-bPAC^m^. Each point represents a cell. Dashed lines indicate 5^th^ and 50^th^ percentiles of GFP-bPAC^m^ expression. Cells that overexpressed bPAC^m^ had higher basal PF fluorescence. This observation alone is consistent with elevated basal cAMP, but could also be explained by co-variation in the expression of bPAC^m^ and PF. **B)** PF (ΔF/F)_max_ upon exposure to a 0.5 second pulse of 488 nm light (234 µJ/mm^2^). Solid curve is an analytical expression (**Supplemental Materials Section 2**) that predicts a bell-shaped curve for PF ΔF/F, assuming that bPAC^m^ has dark activity proportional to its expression level. **C)** PF^m^ ΔF/F response of HEK cells (*n =* 4,005 cells) expressing PF^m^+bPAC^m^ subject to 100 µM forskolin. The decrease in PF^m^ response at high GFP corresponds to the right half of the curve in panel (**B**), indicating that upon overexpression, bPAC dark-state activity elevates basal cAMP. **D)** Same as **(C)**, but for cells expressing only PF^m^ (*n =* 11,334 cells), indicating that PF^m^ sensitivity is independent of expression level.

**Figure S7.**
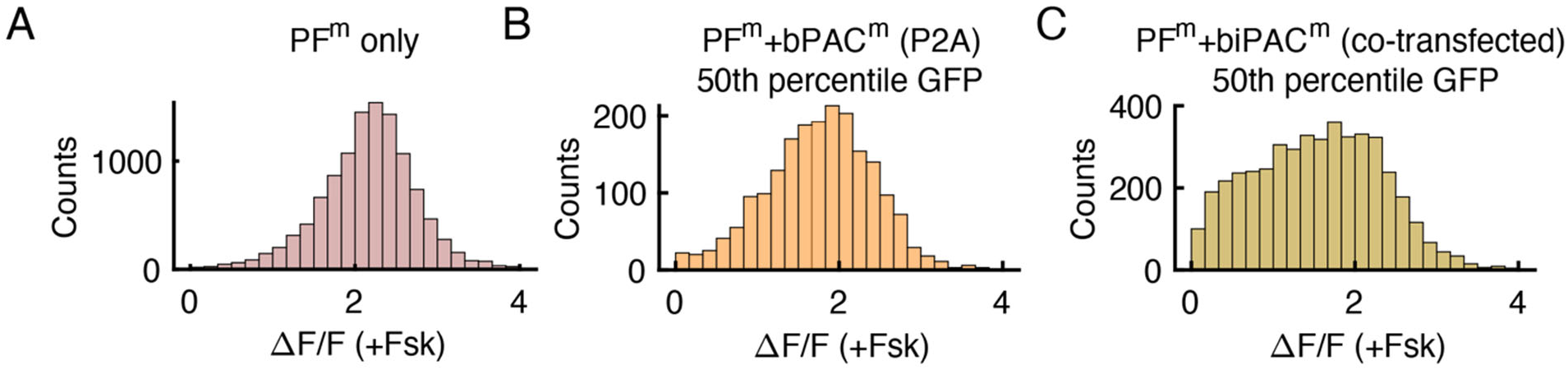
Dark-state activity of bPAC vs. biPAC. **A)** Histogram of ΔF/F for HEK293T cells expressing PF^m^, upon 100 µM Fsk stimulation. Data are from Fig. S6D. Median ΔF/F = 2.2, *n =*11,334 cells. **B)** Same as **A)**, but for cells expressing PF^m^+bPAC^m^. Cells were selected to have < 50^th^ percentile of GFP expression. Median ΔF/F = 1.8, *n* = 2,002 cells. Data are from Fig. S6C. The distribution of forskolin-induced ΔF/F_max_ for cells below the 50^th^ percentile of bPAC^m^ expression largely overlapped with that of cells expressing PF^m^ only. **C)** Same as **B)**, but for cells co-expressing PF^m^ and GFP-biPAC^m^. Median ΔF/F = 1.6, *n* = 4,569 cells.

**Figure S8.**
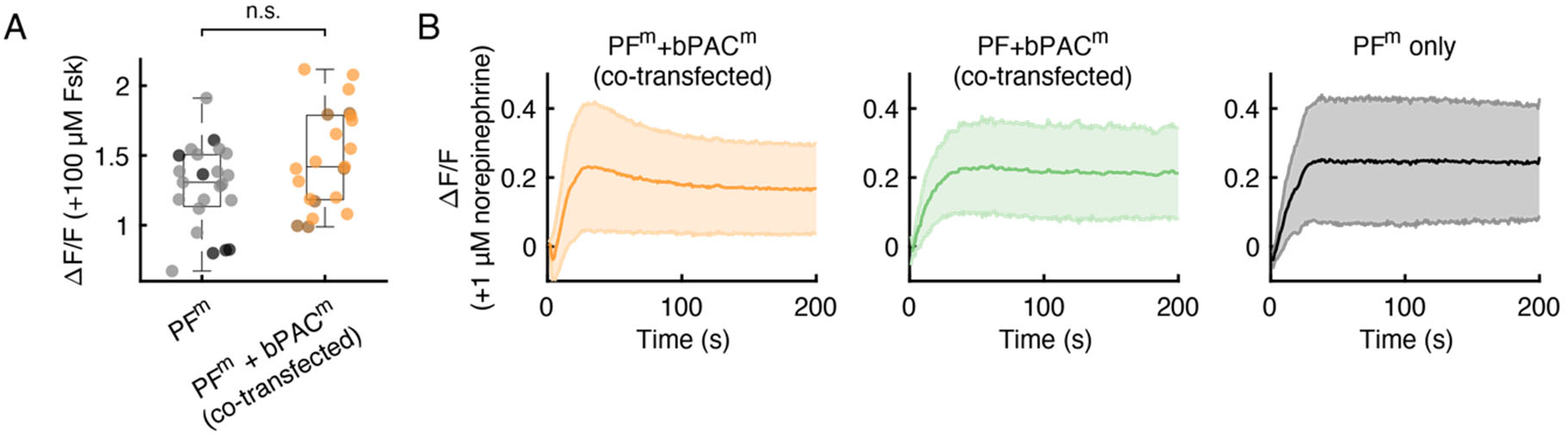
Neuronal bPAC expression does not affect Pink Flamindo responses to either saturating or physiological pharmacological stimuli. **A)** PF ΔF/F upon 100 µM forskolin application for neurons transduced with either only PF^m^ or both PF^m^ and bPAC^m^ (*n =* 23 and 21 neurons respectively, pooled from two experiments each). Boxes represent the interquartile range with the median denoted by the central line. Distributions are not significantly different (*P* = 0.07, Wilcoxon rank-sum test). **B)** Neurons transduced with either PF-bPAC^m^ or with PF^m^ only, responded to 1 µM norepinephrine with ΔF/F ≈ 0.2. Lines represent the mean responses of *n* = 12, 14, and 6 neurons, from left to right. Shading represents s.d..

**Figure S9.**
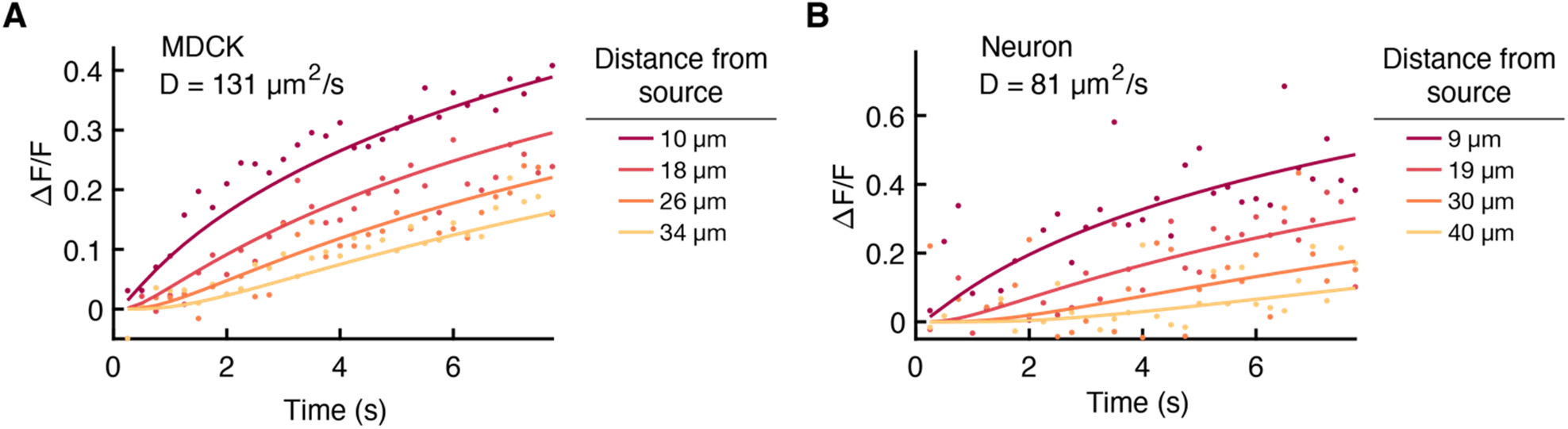
Fitting diffusion coefficients for cells expressing PF^m^+bPAC^m^. **A)** Kymograph traces and fits for the diffusion coefficient for the MDCK cell in Figure 3B. The first 8 s after a 0.5 s pulse of 488 nm light are fit to a 1-dimensional diffusion model. bPAC rate is fixed to be *k*^bPAC^ = 0.05 *s*^−1^. The diffusion coefficient and a scale factor are floated. **B)** Same as **A)**, but for the neuron in Fig. 3D.

**Figure S10.**
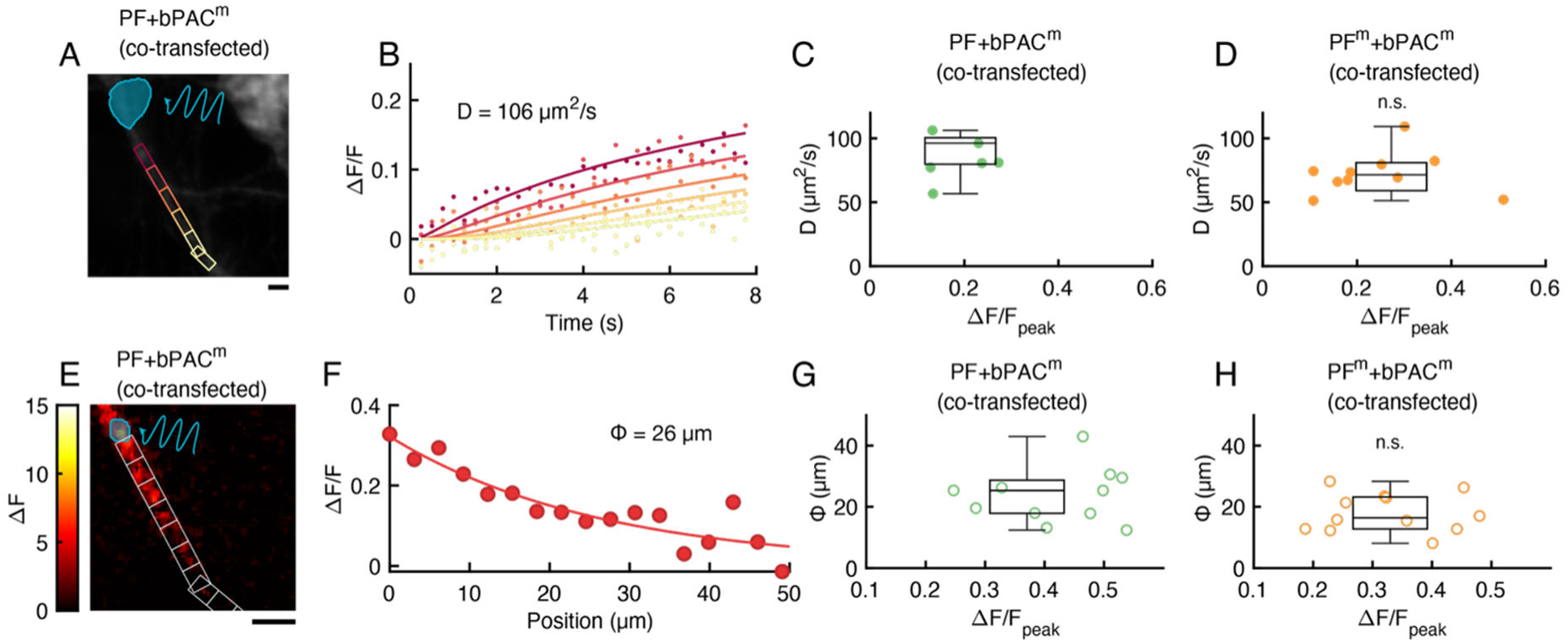
cAMP diffusion coefficient and Thiele length for neurons co-transfected with Pink Flamindo and membrane bPAC. **A)** Image of initial PF fluorescence in a neurite co-expressing PF and GFP-bPAC^m^. Scale bar: 5 µm. **B)** ΔF/F after a 1 s pulse of blue light vs. time, in the indicated regions. Model curves are overlaid, using the best-fit diffusion coefficient. **C)** cAMP diffusion coefficient vs. maximum ΔF/F. *D* = 83 ± 17 µm^2^/s, *n* = 6 neurites from *n* = 3 cells. Blue light intensity was 255 µW/mm^2^. **D)** Same as **C)**, but for cells co-transfected with PF^m^ and bPAC^m^. *D* = 73 ± 17 µm^2^/s, *n* = 10 neurites from *n* = 8 cells. Blue light intensity ranged from 390–1143 µW/mm^2^. All parameters reported as mean ± s.d. Boxes: interquartile range; central line: median. Outliers are excluded. Diffusion coefficients of cells expressing PF+bPAC^m^ were not significantly different compared to cells expressing PF^m^+bPAC^m^ (*P* = 0.18, Wilcoxon rank-sum test). **E)** Average ΔF image of Pink Flamindo fluorescence in a neurite co-expressing PF and GFP-bPAC^m^. 488 nm light was targeted to the blue circled region at an effective optical dose of 51 µW/mm^2^ (1 s of 255 µW/mm^2^ applied every 5 s). Scale bar: 5 µm. **F)** Steady state ΔF/F vs. distance from the targeted region; an exponential fit yielded a cAMP length-scale of 26 µm. **G)** cAMP length-scale vs. ΔF/F close to the blue light-targeted region. ϕ = 24 ± 9 µm, *n* = 11 measurements. Effective blue light intensities ranged from 30–106 µW/mm^2^. **H)** Same as **(G)**, but for neurons co-transfected with PF^m^ and bPAC^m^. ϕ = 18 ± 6 µm, *n* = 12 measurements. Effective blue light intensities ranged from 30– 165 µW/mm^2^. Measured Thiele length for cells expressing PF+bPAC^m^ was not significantly different from cells co-transfected with PF^m^+bPAC^m^ (*P* = 0.10, Wilcoxon rank-sum test). Diffusion coefficients and length constants measured in these experiments were modestly smaller than in Main Text Figs. 3 and 5.

**Figure S11.**
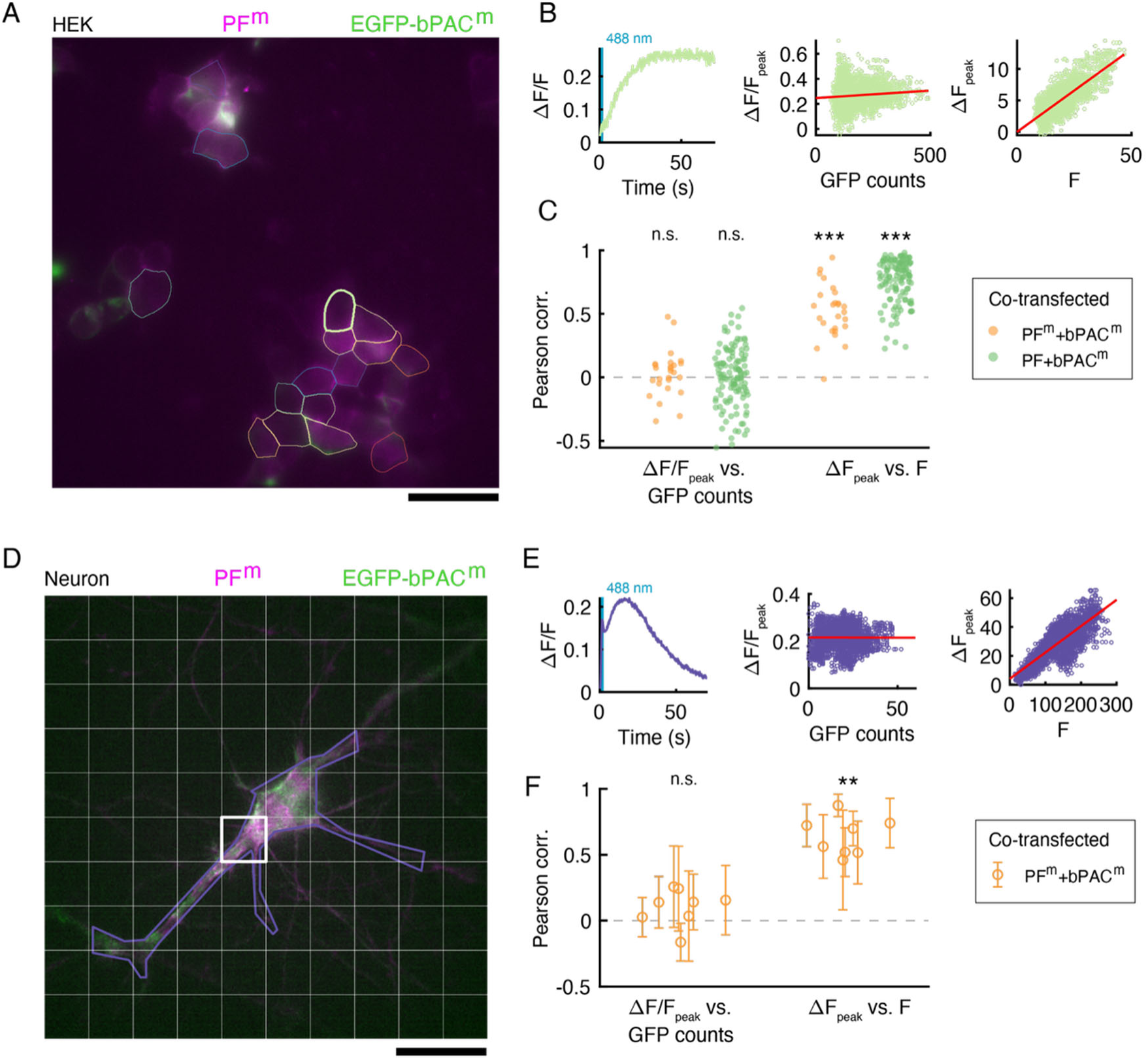
Pixel-wise ΔF/F upon wide-area blue light stimulation. **A)** Image of HEK293T cells co-transfected with PF^m^ (purple) and GFP-bPAC^m^ (green). Scale bar: 50 µm. **B)** Cells were illuminated with a wide area dim pulse of 488 nm light (8.6 –30 µJ/mm^2^). Left to right: (ΔF/F)_peak_ vs. time for the cell indicated in light green from panel **A**, pixel-wise (ΔF/F)_peak_ vs. GFP fluorescence, and pixel-wise ΔF_peak_ vs. F (from PF), for the same cell. Red lines denote the best-fit linear model. **C)** Pearson correlation of (ΔF/F)_peak_ vs. GFP, and of ΔF_peak_ vs. F. Each point represents a cell. Data are shown for cells co-transfected with either PF or PF^m^, and GFP-bPAC^m^. ΔF_peak_ and F are highly correlated, while (ΔF/F)_peak_ and GFP-bPAC are uncorrelated, with the median of the distribution not significantly different from zero. (Left to right: *P* = 0.38, 0.71, 9.3 × 10^-6^, 1.4 × 10^-21^, *n* = 26 cells for PF^m^, *n* = 121 cells for PF, Wilcoxon signed-rank test). **D**–**E**) Same, but for a cultured rat hippocampal neuron. For analysis, the neuron was divided into regions in a 25 by 25 micron square grid, to avoid making correlations between regions much larger than the Thiele length. Scale bar: 50 µm. **F**) Same as **(C),** but each point represents the average across grid regions for a single neuron. Error bars indicate s.d. (Left to right: *P* = 0.11, 0.0078. *n* = 8 cells, Wilcoxon signed-rank test).

**Figure S12.**
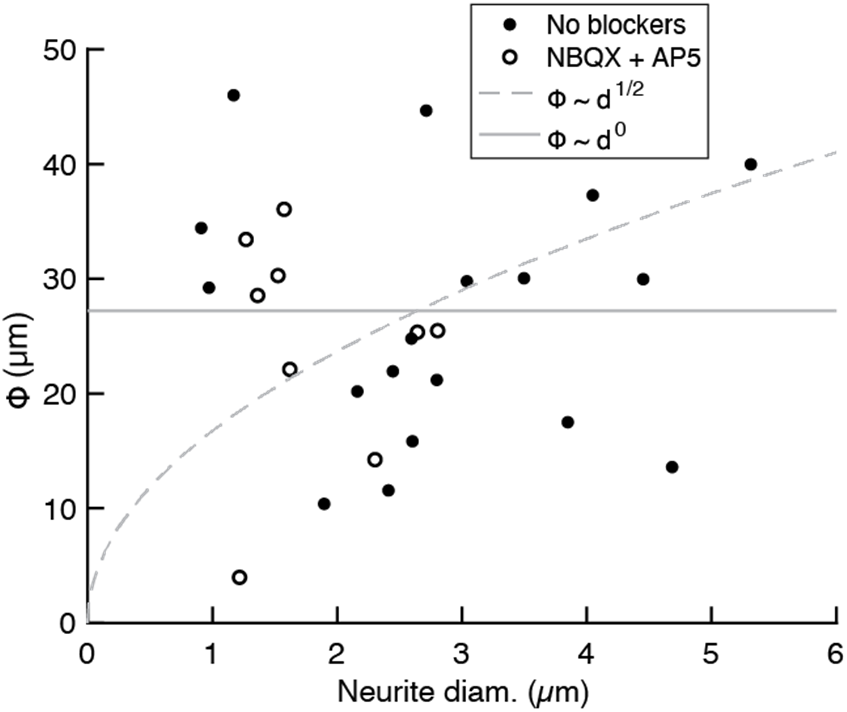
cAMP Thiele length does not follow simple scaling laws implied by soluble or membrane-bound degradation. Thiele length, ϕ, plotted against neurite diameter shows a large variability between neurites, both with and without synaptic blockers (10 µM NBQX, 50 µM D-APV). Lines show theoretical predictions for purely soluble PDE activity (ф ∼ *d*^0^) and purely membrane-bound PDE activity (ф ∼ *d*^1/2^), where *d* is the neurite diameter. The data without blockers are repeated from Fig. 5F.

## Notes

1 If [cAMP] ≥ *K*_M_ then the signal spreads further, so the expression 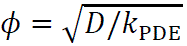 is a lower bound on the signaling length.

## Notes

### Summary of Updates

Control experiments to test that cAMP concentrations are within the physiological range; tests for micron-scale deviations in cAMP concentration around local clusters of bPAC molecules. Updated Introduction and Discussion.

