## Supplementary modeling and calculations for "All-optical mapping of cAMP transport reveals rules of sub-cellular localization"

### Quantitative modeling of cAMP kinetics and diffusion

#### 1 Requirements for nanoscale cAMP compartmentalization

A simple estimate suggests that a single-molecule adenylyl cyclase (AC) source or phosphodiesterase (PDE) sink cannot substantially change the local cAMP concentration. The calculation for an AC source places a maximal bound of +110 nM at the surface of a single AC. The same calculation applies for a PDE sink with the replacement of $k_{\text{cat}}^{\text{AC}}$ by $k_{\text{cat}}^{\text{PDE}}$ and a sign change. A more thorough analysis of this problem is given in Ref. (*1*).

Assuming a finite concentration of PDEs in solution and a single spherical AC source of radius $r_0$, the concentration profile at radii $r > r_0$ is:

$$\delta c(r) = \delta[\text{cAMP}]_0 \frac{r_0}{r} e^{-\frac{r-r_0}{\Phi}}, \tag{1}$$

where $\delta c$ is the deviation from bulk cAMP concentration, $\delta[\text{cAMP}]_0$ is the deviation at the surface of the AC, and $\Phi = \sqrt{D/k_{\text{PDE}}}$ is the Thiele length, the characteristic distance a molecule diffuses before it is degraded. $D$ is the diffusion coefficient and $k_{\text{PDE}} = \frac{k_{\text{cat}}^{\text{PDE}}}{K_m^{\text{PDE}}+[\text{cAMP}]}[\text{PDE}]$ is the PDE reaction rate.

For $\Phi \gg r_0$, the PDEs in solution have little effect on the concentration profile: dilution by diffusion decreases the cAMP concentration before the PDEs can act. For $\Phi \ll r_0$, the concentration profile is essentially the same as for an absorbing or emitting plane. An enzyme radius is $r_0 < 1$ nm while, for realistic PDE concentrations and diffusion coefficients, Thiele lengths are typically tens of microns. Hence $\Phi \gg r_0$.

The total flux from the surface is $J = -4\pi r_0^2 D \frac{\partial c}{\partial r}|_{r=r_0}$, which evaluates to:

$$J = 4\pi\delta[\text{cAMP}]_0 D r_0^2 \left(\frac{1}{r_0} + \frac{1}{\Phi}\right) \tag{2}$$

The maximum deviation in concentration at the enzyme surface occurs as $\Phi \to \infty$, i.e. in the absence of cAMP degradation in the bulk. In this case, $\delta[\text{cAMP}]_0 = J/(4\pi D r_0)$. The maximal flux is set by the turnover rate of the enzyme, i.e. $J_{max} = k_{\text{cat}}^{\text{AC}}$ or $k_{\text{cat}}^{\text{PDE}}$. Myocardial adenylyl cyclases have a turnover of 40 $s^{-1}$ and olfactory cilia adenylyl cyclases have a turnover of 140 $s^{-1}$ (*2*). The turnover of bPAC in the light is 2.6 ± 0.3 $s^{-1}$ (*3*). The literature values for $k_{\text{cat}}^{\text{PDE}}$ range from 5 $s^{-1}$ to 20 $s^{-1}$ (*4–7*). Assuming an effective enzyme radius of $r_0 = 1$ nm, the maximal deviations in concentration at the surface of a single endogenous AC enzyme are $\delta[\text{cAMP}]_0 = +30$ to $+110$ nM (and even smaller for bPAC), and at the surface of a single PDE enzyme are $\delta[\text{cAMP}]_0 = -4$ to $-15$ nM. Since basal concentrations of cAMP in cells range from 1 - 5 µM (*8–11*), these local deviations in concentration are likely insignificant. The prospect for clustering of enzymes to induce localized deviations in cAMP concentration has been analyzed in detail in Ref. (*1*).

One can also ask at what concentration of PDE the Thiele length becomes comparable to the molecular size, $r_0$. For a PDE $k_{\text{cat}}^{\text{PDE}}$ of 10 $s^{-1}$, $D = 120$ μm$^2/s$, and $K_m = 1$ μM (*10*) and [cAMP] = 1 μM, a PDE concentration of 5 M would be required for $\Phi$ to reach 2 nm.

### 2 Modeling of bPAC dark-state activity

To quantiy bPAC dark-state activity, we seek an expression for Pink Flamindo $\Delta F/F$ as a function of bPAC expression level, [bPAC]. We assume that bPAC has some dark state activity proportional to its expression level, and that it has bright state activity proportional to both its expression level and the blue light intensity. Then:

$$[\text{cAMP}]_0 = c_{\text{basal}} + \alpha[\text{bPAC}]$$
$$[\text{cAMP}]_f = c_{\text{basal}} + (\alpha + I\beta)[\text{bPAC}]$$

where $\alpha$ is a parameter associated with cAMP production due to bPAC dark-state activity, and $\beta$ is associated with cAMP production due to activated bPAC in the light.

Pink Flamindo $\Delta F/F$ follows:

$$\Delta F/F = \frac{H([\text{cAMP}]_f) - H([\text{cAMP}]_0)}{H([\text{cAMP}]_0)}, \tag{3}$$

where $[\text{cAMP}]_0$ and $[\text{cAMP}]_f$ are the cAMP concentrations before and after a given stimulus, and $H([\text{cAMP}])$ is the Hill equation: $H([\text{cAMP}]) \equiv \frac{[\text{cAMP}]^n}{K_D + [\text{cAMP}]^n}$, where $K_D = 7.2\ \mu$M is the Pink Flamindo dissociation constant (*12*) and [cAMP] is the cAMP concentration. $n = 1.01$ is the Hill coefficient of Pink Flamindo.

In Figure S6, we used GFP counts as a proxy for [bPAC]. Figure S6B shows Equation 3, with $c_{\text{basal}} = 3\ \mu$M, $\alpha = 10^{-3}\ \mu$M·(GFP counts)$^{-1}$, $I\beta = 0.1\ \mu$M·(GFP counts)$^{-1}$ at a near-saturating intensity of $I = 234\ \mu$J/mm$^2$ of 488 nm light.

### 3 Modeling of cAMP dynamics

When bPAC receives a brief pulse of 488 nm light, it converts to the active state and produces cAMP. The activated bPAC decays back to the inactive state.

$$\text{bPAC} \underset{k_{\text{rev}}^{\text{bPAC}}}{\overset{Ik_E}{\rightleftharpoons}} \text{bPAC*}$$

bPAC: inactivated bPAC

bPAC*: activated bPAC

$I$: 488 nm laser intensity

$k_E$: bPAC formation rate constant

$k_{\text{rev}}^{\text{bPAC}}$: bPAC relaxation rate constant

We denote the concentration of bPAC* as [AC(x,t)]. The concentration of cAMP then depends on the amount of activated bPAC as well as the activity of phosphodiesterases (PDEs).

$$\text{ATP} \xrightarrow{k_{\text{cat}}^{\text{AC}} \cdot [\text{AC(x, t)}]} \text{cAMP} + \text{PP}_i \xrightarrow{k_{\text{PDE}}} \text{AMP}$$

$k_{\text{cat}}^{\text{AC}}$: bPAC turnover rate
[AC(x, t)]: concentration of activated bPAC
$k_{\text{PDE}}$: rate of cAMP degradation by PDEs

The effective PDE rate constant, $k_{\text{PDE}}$, also depends on the concentration of PDEs. $k_{\text{PDE}} = \frac{k_{\text{cat}}^{\text{PDE}}}{K_m^{\text{PDE}} + [\text{cAMP}]}[\text{PDE}] \approx \frac{k_{\text{cat}}^{\text{PDE}}}{K_m^{\text{PDE}}}[\text{PDE}]$ when $[\text{cAMP}] \ll K_m^{\text{PDE}}$, where $k_{\text{cat}}^{\text{PDE}}$ is the PDE turnover rate, and $K_m^{\text{PDE}}$ is the PDE Michaelis constant.

We assume that ATP is in excess and that degradation is first order in cAMP concentration and in PDE concentration. We assume that the PDE concentration does not change, since it is an enzyme. Finally, cAMP binding to Pink Flamindo is faster than these timescales and reversible, hence we do not include it in the model. Then, we have:

$$\frac{\partial c}{\partial t} = D\nabla^2 c - k_{\text{PDE}} c + k_{\text{cat}}^{\text{AC}} \cdot [\text{AC(x, t)}] \tag{4}$$

where $c$ is the cAMP concentration and $D$ is the diffusion coefficient. All of the modeling in the paper is based on Equation 4. We consider different AC activation profiles [AC(x,t)], reflecting different patterns of GPCR inputs.

| Fit variable | Description | Units |
|---|---|---|
| $k_{\text{PDE}}$ | PDE reaction rate | $s^{-1}$ |
| $k_{\text{rev}}^{\text{bPAC}}$ | bPAC relaxation rate constant | $s^{-1}$ |
| $D$ | Diffusion coefficient | $\mu m^2/s$ |
| Defined variable | | |
| $k_{\text{cat}}^{\text{AC}}$ | bPAC turnover rate | $s^{-1}$ |
| $k_F$ | bPAC turnover rate × activated bPAC concentration | moles/liter $s^{-1}$ |
| $k_{\text{cat}}^{\text{PDE}}$ | PDE catalytic rate constant | $s^{-1}$ |
| $K_m^{\text{PDE}}$ | PDE Michaelis constant | moles/liter |

Table S1: Table of units.

#### 3.1 Spatially homogeneous transient dynamics

After a wide-area $\delta$-function pulse of blue light, the subsequent evolution of [AC] is: $\frac{d}{dt}[\text{AC}] = -k_{\text{rev}}^{\text{bPAC}}[\text{AC}]$, and $[\text{AC(t)}] = [\text{AC}]_0 e^{-k_{\text{rev}}^{\text{bPAC}} t}$. $[\text{AC}]_0$ is the concentration of active bPAC right after the blue light exposure, which is proportional to the total bPAC concentration as well to the blue light dose. Since the stimulus is spatially homogeneous, we set the diffusive term in Equation 4 to zero, and thus:

$$\begin{aligned}\frac{dc}{dt} &= -k_{\text{PDE}} \cdot c + k_{\text{cat}}^{\text{AC}} \cdot [\text{AC(t)}] \\ &= -k_{\text{PDE}} c + k_{\text{cat}}^{\text{AC}} \cdot [\text{AC}_0] e^{-k_{\text{rev}}^{\text{bPAC}} t}\end{aligned}$$

which is solved by:

$$\Delta\text{cAMP}(t) = k_F \cdot \left( \frac{e^{-k_{\text{PDE}} t} - e^{-k_{\text{rev}}^{\text{bPAC}} t}}{k_{\text{rev}}^{\text{bPAC}} - k_{\text{PDE}}} \right). \tag{5}$$

We defined $k_F \equiv k_{\text{cat}}^{\text{AC}} \cdot [\text{AC}]_0$. The formula is the same under interchange of the two rate constants. We observed experimentally that the PDE degradation rate was slower than the bPAC* relaxation rate, so in the fit we assigned the two fitting constants using this ranking. This expression is plotted and fit to experimental data in Figure 2F.

For Figure 2F, we averaged the peak-normalized single-cell $\Delta F/F$ traces, and fit the average trace for visualization purposes. For each construct, single-cell traces were fit with a shared $k_{\text{rev}}^{\text{bPAC}}$ across all cells but a $k_{\text{PDE}}$ specific for each cell. Our logic was that $k_{\text{rev}}^{\text{bPAC}}$ is an intensive property of the bPAC, unlikely to vary between cells, while $k_{\text{PDE}}$ depends implicitly on PDE concentration and activity, which might vary substantially between cells.

### 4 1D cAMP diffusion

#### 4.1 cAMP spread from a transiently active localized source

We consider cAMP diffusion in a 1D tube where the diameter of the tube, $d$, is much smaller than the length, $L$, and the space is half-infinite with a reflecting boundary at $x = 0$. The general solution for the change from baseline in cAMP concentration, which we denote as $c(x, t)$, is given by:

$$c(x, t) = \int_0^t \int_0^x \left( k_{\text{cat}}^{\text{AC}} \cdot [\text{AC}(x', t')] \cdot G(x - x', t - t') \right) dx' dt' \tag{6}$$

Where $G(x, t)$ is the Green's function for Equation 4:

$$G(x, t) = \underbrace{\frac{e^{-\frac{x^2}{4Dt}}}{\sqrt{\pi D t}}}_{\text{diffusive spreading}} \cdot \underbrace{e^{-k_{\text{PDE}} t}}_{\text{degradation by PDEs}}$$

We consider AC activation by a pulse of blue light that is a $\delta$-function in time and is localized between $x = 0$ and $x = \Delta x$, where $\Delta x$ is small compared to the length scales of interest. Thus, [AC(x,t)] is a decaying exponential in time, as above. For this case, we have:

$$c(x, t) = \int_0^t \int_0^{\Delta x} \left( k_F e^{-k_{\text{rev}}^{\text{bPAC}} t'} \cdot \underbrace{\frac{e^{-\frac{(x-x')^2}{4D(t-t')}}}{\sqrt{\pi D (t - t')}}}_{\text{diffusive spreading}} \cdot \underbrace{e^{-k_{\text{PDE}}(t-t')}}_{\text{degradation by PDEs}} \right) dx' dt'$$

We perform the integral over $x'$ first and get:

$$c(x,t) = \int_0^t \left( \frac{k_F e^{-k_{\text{rev}}^{\text{bPAC}} t'}}{\sqrt{\pi D(t-t')}} \cdot e^{-k_{\text{PDE}}(t-t')} \cdot \frac{\sqrt{4\pi D(t-t')}}{2} \left( \text{erf}\left( \frac{\Delta x - x}{\sqrt{4D(t-t')}} \right) + \text{erf}\left( \frac{x}{\sqrt{4D(t-t')}} \right) \right) \right) dt'$$
$$\approx \int_0^t \left( k_F \Delta x \cdot e^{-k_{\text{rev}}^{\text{bPAC}} t'} \cdot \frac{e^{-\frac{x^2}{4D(t-t')}}}{\sqrt{\pi D(t-t')}} \cdot e^{-k_{\text{PDE}}(t-t')} \right) dt'$$

We assumed that $\Delta x$ is small and expanded to first order in $\Delta x$. Performing the integral over $t'$:

$$c(x,t) = \frac{k_F \Delta x \cdot e^{-\frac{x\sqrt{k_{\text{rev}}^{\text{bPAC}} - k_{\text{PDE}}}}{\sqrt{D}} - k_{\text{PDE}} t} \left( \text{erfc}\left( \frac{\frac{x}{\sqrt{D}} - 2t\sqrt{k_{\text{rev}}^{\text{bPAC}} - k_{\text{PDE}}}}{2\sqrt{t}} \right) - e^{\frac{2x\sqrt{k_{\text{rev}}^{\text{bPAC}} - k_{\text{PDE}}}}{\sqrt{D}}} \text{erfc}\left( \frac{\frac{x}{\sqrt{D}} + 2t\sqrt{k_{\text{rev}}^{\text{bPAC}} - k_{\text{PDE}}}}{2\sqrt{t}} \right) \right)}{2\sqrt{D}\sqrt{k_{\text{rev}}^{\text{bPAC}} - k_{\text{PDE}}}} \quad (7)$$

If the point source is not next to a reflecting boundary but rather at the midpoint of the tube, the expression has an additional factor of 2 in the denominator.

##### 4.2 Analysis

Data were fit in MATLAB to Equation 7 above. We decided to fit only short times to obtain a more accurate measurement of $D$. We fit data from times $t < 8$ s and set $k_{\text{PDE}} = 0$ since it was ill-constrained for short times. For each cell, we computed the background-subtracted $\Delta F/F$ time traces for several regions along the long axis of the cell. We performed a fit across the data pooled from these traces. We fixed $k_{\text{rev}}^{\text{bPAC}} = 0.05\ \text{s}^{-1}$ based on whole-cell measurements, and also fixed $x$, the distance from the source, for each kymograph trace. We floated only the diffusion coefficient $D$.

#### 5 cAMP length-scale in neurons

##### 5.1 cAMP spread from a continuously active localized source

We consider Equation 4 in an infinite 1D tube, with a spatially localized source of width $\Delta x$, centered at $x = 0$. At steady state, $\frac{\partial c}{\partial t} = 0$, and the equation is solved by the decaying exponential

$$c(x) = [\text{cAMP}]_0 e^{-|x|/\Phi} \quad (8)$$

where $[\text{cAMP}]_0 = \frac{k_F \Delta x}{2\sqrt{D k_{\text{PDE}}}}$ is a prefactor and $\Phi \equiv \sqrt{D/k_{\text{PDE}}}$ is the characteristic length scale over which the cAMP concentration decreases by a factor of $e$.

To compute the prefactor, we consider the concentration at positive values of $x$, and take the derivative $\frac{\partial c}{\partial x} = -c(x)/\Phi$ and note that the flux $-D\frac{\partial c}{\partial x} = \frac{1}{2} k_F \Delta x$ (units: molecules cAMP / length$^2$; half the flux goes to positive $x$ and half to negative $x$). Then $c(x) = \frac{1}{2}(\Phi k_F \Delta x / D) e^{-x/\Phi} = \frac{k_F \Delta x}{2\sqrt{D k_{\text{PDE}}}} e^{-x/\Phi}$ which has units of cAMP concentration.

For completeness, we also compute the full concentration profile in time and space for a localized source that turns on at $t = 0$ and remains continually active, using the same strategy as in Section 4.1:

$$\begin{aligned} c(x,t) &= \int_0^t \left( k_F \Delta x \frac{e^{-\frac{x^2}{4D(t-t')}}}{2\sqrt{\pi D(t-t')}} \cdot e^{-k_{\text{PDE}}(t-t')} \right) dt' \\ &= \int_0^t \left( k_F \Delta x \frac{e^{-\frac{x^2}{4Dt'}}}{2\sqrt{\pi D t'}} \cdot e^{-k_{\text{PDE}} t'} \right) dt' \end{aligned}$$

which we solve in Mathematica to get:

$$c(x,t) = \frac{k_F \Delta x \cdot e^{-\frac{x\sqrt{k_{\text{PDE}}}}{\sqrt{D}}} \left( \text{erfc}\left( \frac{\frac{x}{\sqrt{D}} - 2t\sqrt{k_{\text{PDE}}}}{2\sqrt{t}} \right) - e^{\frac{2x\sqrt{k_{\text{PDE}}}}{\sqrt{D}}} \text{erfc}\left( \frac{\frac{x}{\sqrt{D}} + 2t\sqrt{k_{\text{PDE}}}}{2\sqrt{t}} \right) \right)}{4\sqrt{D k_{\text{PDE}}}}$$

At the center, $c(0,t) = k_F \Delta x \cdot \text{erf}(\sqrt{k_{\text{PDE}} t})/(2\sqrt{D k_{\text{PDE}}})$ which at steady-state ($t \to \infty$) goes to $k_F \Delta x/(2\sqrt{D k_{\text{PDE}}})$ in agreement with above.

If the point source is at the closed end of a semi-infinite tube, then all the concentrations in this section should be multiplied by 2.

#### 5.2 Analysis

We fit our data to Equation 8 with free parameters $[\text{cAMP}]_0$ and $\Phi$. Steady-state average $\Delta F/F$ images were 2D median filtered over space with a neighborhood size of 4 pixels for visualization purposes. Only frames without the blue illumination were included. Length scale analysis data were taken when $\text{PF}^{\text{m}}$ $\Delta F/F$ reached steady state. Data were median filtered over time with a filter size of 40 frames, and background subtracted according to the 35th percentile fluorescence of pixels in each background region to either side of the region along the neurite being analyzed. Data on each side of the illumination region were fit separately. We avoided including neurites with significant sub-branching structures.

Neurite widths were computed by measuring the full width half maximum of a Gaussian fit to the time-averaged background-subtracted fluorescence, across a line section cut perpendicular to the long axis of each neurite with a bin size of 0.5 by 0.5 microns.

### 6 Scaling of concentrations and timescale with system size

To build intuition for how geometry and enzyme localization bias second-messenger signaling, we consider a minimal reaction–diffusion model in a tube-shaped cellular process with diameter $d$ and length $L$. We focus on axial length scales $|x| \gtrsim d$ where transverse gradients can be neglected, so that $c(x,t)$ is effectively one-dimensional. Starting from Eq. 4, the corresponding 1D model is

$$\frac{\partial c}{\partial t} = D \frac{\partial^2 c}{\partial x^2} - k_{\text{PDE}}\, c + k_F(x,t), \tag{9}$$

where $k_F(x,t) = k_{\text{cat}}^{\text{AC}}[\text{AC}(x,t)]$ is the local cAMP production rate (set by the concentration of active adenylyl cyclase), and $k_{\text{PDE}}$ is the effective first-order degradation rate (set by the concentration of active PDE in the approximately linear regime of degradation).

A key geometric point is that the *effective volumetric concentration* of a membrane-bound enzyme scales with surface-to-volume ratio, whereas a concentration of a soluble enzyme does not. For both a tube and sphere, $S/V \propto 1/d$. Thus if an enzyme is membrane-localized with areal density $\sigma$ (molecules per membrane area), its effective volumetric concentration scales as $[\text{enzyme}]_{\text{eff}} \propto \sigma\,(S/V) \propto 1/d$, while for soluble enzymes $[\text{enzyme}]_{\text{eff}} \propto d^0$. The scaling relations below follow directly from these geometric conversions together with Eq. 9. These ideas are also explored in Ref. (*13*).

#### 6.1 Homogeneous activation: Concentration scaling

We solve Equation 9 with $D\frac{\partial^2 c}{\partial x^2} = 0$, $\frac{\partial c}{\partial t} = 0$, and $k_\text{F}$ constant everywhere. Then $c = k_\text{F}/k_{\text{PDE}} \propto \frac{[\text{AC}]}{[\text{PDE}]}$ and therefore if both bPAC and PDEs are soluble, $c \propto d^0/d^0 = d^0$, and if both are membrane-bound, $c \propto d^{-1}/d^{-1} = d^0$.

For soluble PDE and membrane-bound AC, $c \propto d^{-1}$.

For membrane-bound PDE and soluble AC, $c \propto d$.

#### 6.2 Homogeneous activation: Time constant scaling

For homogeneous activation, the time constant to reach steady-state can be determined from the solution to Equation 9 with the diffusive term set to zero and with the initial condition $c(0) = 0$. The solution is $c(t) = \frac{k_\text{F}(1-e^{-k_{\text{PDE}}t})}{k_{\text{PDE}}}$, and therefore the time constant $\tau = 1/k_{\text{PDE}}$. For soluble PDE, $\tau \propto d^0$, and for membrane-bound PDE, $\tau \propto d$.

#### 6.3 Point-source activation: Thiele length scaling

For a localized, stationary source centered at $x = 0$ with small axial extent $\Delta x$ (idealized as $k_F(x) = k_F\,\Delta x\,\delta(x)$), the steady-state solution to Eq. 9 is an exponential profile,

$$c(x) = c_{\text{bg}} + [\text{cAMP}]_0\, e^{-|x|/\Phi}, \tag{10}$$

where $c_{\text{bg}}$ is the background concentration far from the source, $[\text{cAMP}]_0$ is the peak concentration at the source, and

$$\Phi = \sqrt{D/k_{\text{PDE}}} \tag{11}$$

is the Thiele length (the characteristic distance a molecule diffuses before being degraded).

$\Phi$ depends on the *localization of the sink* (PDE) but not on the localization of the source (AC), because diffusion and degradation set the decay away from the source. Specifically, $\Phi \propto \sqrt{1/[\text{PDE}]}$. Thus for soluble PDEs, $\Phi \propto d^0$, and for membrane-bound PDEs, $\Phi \propto \sqrt{d}$.

#### 6.4 Point-source activation: Prefactor scaling

In the linear regime, $[\text{cAMP}]_0$ is set by the balance of localized production with axial diffusion and degradation. The peak concentration for point-source activation is $[\text{cAMP}]_0 = \frac{k_F \Delta x}{2\sqrt{D k_{\text{PDE}}}}$. Thus if both PDE and AC are soluble, both the numerator $k_F \propto d^0$ and denominator $k_{\text{PDE}} \propto d^0$ so $[\text{cAMP}]_0$ is independent of $d$.

For soluble PDE but membrane-bound AC, $[\text{cAMP}]_0 \propto \frac{1/d}{\sqrt{d^0}} = d^{-1}$.

For membrane-bound PDE but soluble AC, $[\text{cAMP}]_0 \propto \frac{d^0}{\sqrt{1/d}} = d^{1/2}$.

Finally, if both PDE and AC are membrane-bound, $[\text{cAMP}]_0 \propto \frac{1/d}{\sqrt{1/d}} = d^{-1/2}$.

A corollary of the above scaling rules is that the spatially integrated excess signal for a point source,

$$\int_{-\infty}^{\infty} (c(x) - c_{\text{bg}})\, dx = 2\,[\text{cAMP}]_0\,\Phi, \tag{12}$$

scales like $1/k_{\text{PDE}}$ because $[\text{cAMP}]_0 \propto 1/\sqrt{k_{\text{PDE}}}$ and $\Phi \propto 1/\sqrt{k_{\text{PDE}}}$. Thus, up to the conversion between a localized production rate ($k_F \Delta x$) and a distributed production rate ($k_F$), the integrated excess concentration has the same overall $k_{\text{PDE}}$-dependence (and hence the same overall diameter scaling) as the homogeneous-production steady state.
